# Trajectory Analysis of Hepatic Stellate Cell Differentiation Reveals Metabolic Regulation of Cell Commitment and Fibrosis

**DOI:** 10.1101/2024.02.15.577777

**Authors:** Raquel A. Martínez García de la Torre, Julia Vallverdú, Qing Xu, Silvia Ariño, Beatriz Aguilar-Bravo, Paloma Ruiz-Blázquez, Maria Fernandez-Fernandez, Artur Navarro-Gascon, Albert Blasco-Roset, Paula Sànchez-Fernàndez-de-Landa, Juan Pera Garcia, Damià Romero-Moya, Paula Ayuso Garcia, Celia Martínez Sánchez, Laura Zanatto, Laura Sererols, Paula Cantallops Vilà, Bénédicte Antoine, Mikel Azkargorta, Juan José Lozano, Maria L Martínez-Chantar, Alessandra Giorgetti, Félix Elortza, Anna Planavila, Marta Varela, Ashwin Woodhoo, Antonio Zorzano, Isabel Graupera, Anna Moles, Mar Coll, Silvia Affo, Pau Sancho-Bru

## Abstract

Defining the trajectory of cells during differentiation and disease offers the possibility to understand the mechanisms driving cell fate and identity. However, trajectories of human cells are largely unexplored. By investigating the proteome trajectory of iPSCs differentiation to hepatic stellate cells (dHSCs), we identified RORA as a key transcription factor governing the metabolic reprogramming of HSCs necessary for HSCs’ commitment, identity, and activation. Using RORA deficient iPSCs and pharmacologic interventions, we showed that RORA is required for mesoderm differentiation and prevents dHSCs activation by reducing the high energetic state of the cells. While RORA knockout mice had enhanced fibrosis, RORA agonists rescued multi-organ fibrosis in *in vivo* models. RORA expression was consistently found to be negatively correlated with liver fibrosis and HSCs activation markers in patients with liver disease. This study reveals that RORA regulates cell metabolic plasticity, crucial for mesoderm differentiation, pericyte quiescence, and fibrosis, influencing cell commitment and disease mechanisms.

**Summary:** This study describes the trajectory of induced pluripotent stem cells (iPSCs) differentiation to hepatic stellate cells (dHSCs). We identify RAR-related orphan receptor alpha (RORA) as a transcription factor essential for mesoderm commitment and dHSCs identity and fibrogenic activation by regulating metabolic plasticity.

## Introduction

During embryonic development cells undergo a transcriptomic and metabolic reprogramming controlled by pathways tightly regulated in a time, space, and cell dependent manner. When the embryonic development ends, homeostasis takes over. Nonetheless, developmental signaling pathways can be reactivated for the maintenance and repair of adult tissues^1^. Therefore, it is fundamental to investigate the mechanisms governing cell identity and their fate upon development and disease to better define the differentiation paths and understand cell trajectories in space and time. In this context, human induced pluripotent stem cells (iPSCs) are a promising and useful tool, as they can differentiate into any cell type, and can be shaped *in vitro* to mimic a disease condition, offering the possibility to study cell physiological and pathophysiological trajectory ^2,3^.

Fibrosis is an evolutionary conserved mechanism involved in multiple chronic injury conditions and tissues and is responsible for nearly 45% of all death in industrialized countries, being the liver one of the principal organs affected^4^. Liver fibrosis is the common outbreak of most liver diseases and is associated with the risk of mortality and liver-related morbidity^5^. The main cell type responsible for liver fibrosis are hepatic stellate cells (HSCs)^6^. These cells are specialized liver pericytes that maintain extracellular matrix (ECM) homeostasis of the organ and store vitamin A in cytoplasmic lipid droplets^7^. During liver homeostasis, HSCs are in a quiescent state, but in response to injury, they activate and acquire a myofibroblastic-like phenotype, participating in the wound-healing response to injury and in tissue regeneration. In chronic liver injury, activation of HSCs persist and they become the main fibrogenic cell type in the liver ^8,9^. Cell metabolism is crucial for the activation of HSCs, which, to fulfil the high metabolic requirements during activation, increases glycolysis^10^, glutaminolysis^11^ and *de novo* lipogenesis^12^.

The maintenance of the HSCs quiescent phenotype is tightly regulated by transcription factors (TF). Some of these TF are expressed during liver development and become downregulated during HSCs activation^13,1415^. Restoration of their expression in activated HSCs, re-establish, in part, the quiescent state, indicating a parallelism between both processes of maturation and fibrogenesis^13,1415^. Thus suggesting that understanding the trajectory of HSCs in development and disease may help to identify molecular pathways suitable for preventing HSCs activation or promoting the regression to a quiescent phenotype, thus mitigating fibrogenesis in chronic liver diseases.

In this study, we performed a time-resolving proteome characterization of iPSC differentiation to functional HSCs (dHSCs), which mimic phenotypic and functional characteristics of primary human HSCs^6,7^. Furthermore, we identify retinoic acid receptor -related orphan receptor alpha (RORA) as a TF regulating both HSCs differentiation and favouring the maintenance of a quiescent phenotype by modulating the metabolic state of the cells. In both, mesoderm commitment and dHSCs activation downregulation of RORA mediates a metabolic switch, increasing glycolysis and mitochondrial oxidative phosphorylation system (OXPHOS). Moreover, we confirm the anti-fibrogenic role of RORA in hepatic and extrahepatic pericytes, thus positioning RORA as a potential antifibrogenic multiorgan core target. Overall, this work shows the potential of studying cell trajectories along differentiation and disease progression.

## Results

### Time-resolving proteomic analysis of dHSCs differentiation

To have a complete proteome profile of the iPSC differentiation towards functional iPSC-differentiated HSCs (dHSCs), we obtained 7 different timepoints across differentiation (day 0, day 2, day 4, day 6, day 8, day 10 and day 12) as previously described (Coll et al., 2018; Vallverdú et al., 2021)^16,17^. Samples from four independent differentiations were collected and assessed using mass spectrometry (MS)-based profiling (Figure 1A). We identified 3064 proteins, of which 2475 were quantified, providing a comprehensive proteomic roadmap of the differentiation process from iPSCs to dHSCs.

**Figure 1:**
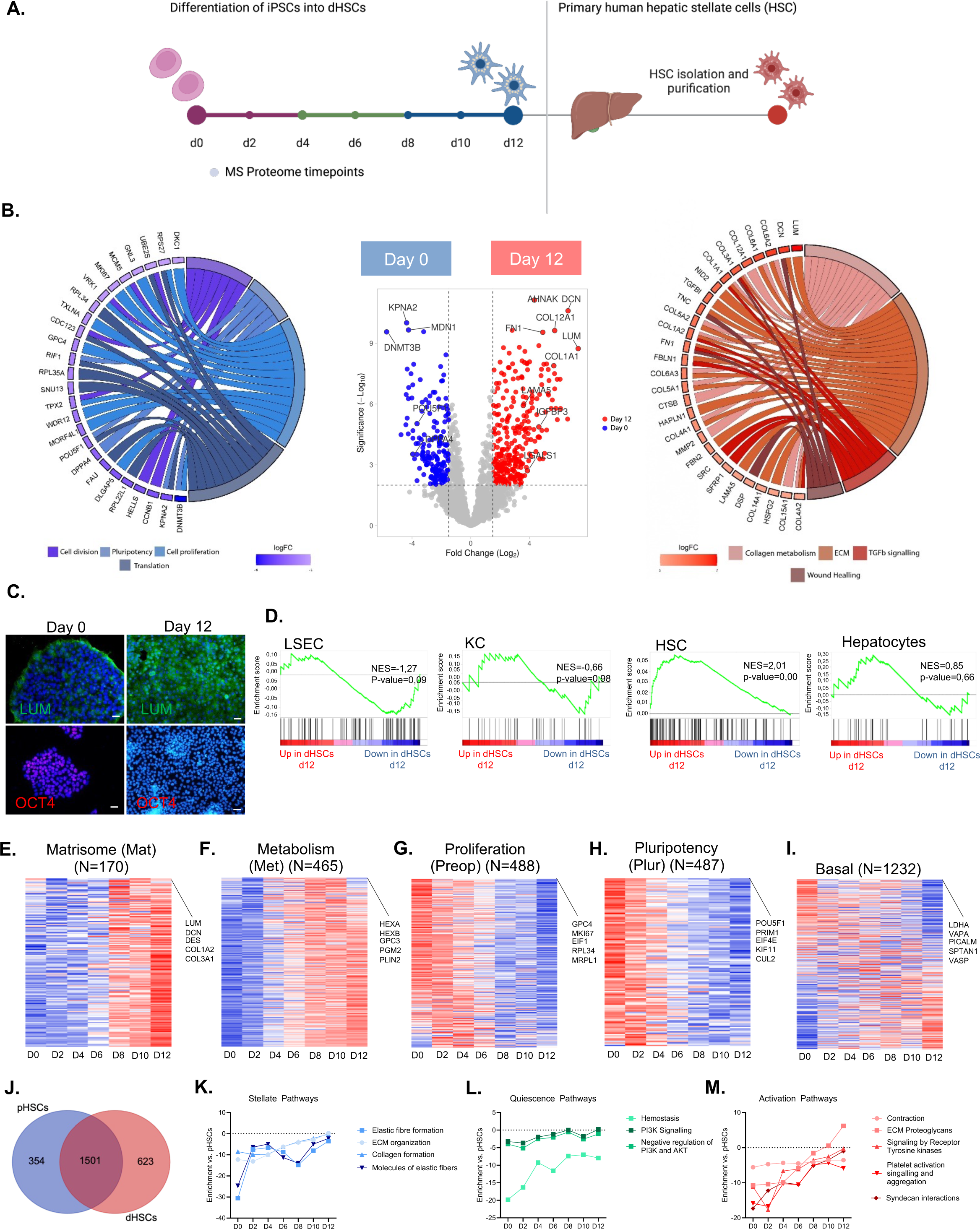
Time-resolved proteomic characterization of the iPSCs differentiation to HSCs shows that the stellate cell phenotype is acquired gradually. (A) Schematic illustration of the experimental design for the characterization at a proteome level of the differentiation process and comparison to primary HSCs. (B) Chord diagram showing the GOs enriched at day 0 and day 12 and volcano plot showing the differentially expressed proteins of the direct comparison between both timepoints. (C) Representative images of immunofluorescence at day 0 and day 12 of LUM (Lumican) and OCT4 (POU Class 5 Homeobox 1). Scale bars at day 2 represent 200μm and at day 12 100μm(D) GSEAs of reported proteomic signatures from human liver cell types (LSEC – Liver sinusoidal epithelial cells; KC – Kupffer Cells; HSC – Hepatic Stellate Cells and HEP – Hepatocytes). (E-I) Protein profile of the 5 identified clusters across differentiation. In graphics, D0 is day 0; D2 is day2; D4 is day 4; D6 is day 6; D8 is day 8; D10 is day 10; d12 is day 12. (J) Venn Diagram illustrating that both cell types have a similar proteome profile. (K-M) dHSCs gained the expression of pathways related to quiescence and activation phenotype, as well as common stellate cell phenotype pathways.

As expected, during the differentiation HSC markers (DCN, LUM, PTN, MMP2, PCOLCE, LAMA5, IGFBP3, LGALS1, IGFBP5, and collagens) increased and pluripotency proteins (RIF1, POU5F1, DPPA4, KPNA2, DNMT3B) decreased (Figure 1B and C). Specifically, day 0 proteins were enriched in cell proliferation, translation, cell division, and pluripotency gene ontologies (GOs). Conversely, at day 12 we observed an enrichment in proteins related to ECM organization, collagen metabolism, wound healing, and TGFβ signaling (Figure 1B; Table S1A-B). Gene Set Enrichment Analysis (GSEA) of proteome signatures of human liver cell types^18^ showed that the dHSC proteome profile was significantly enriched in the HSC signature, resembling the human primary HSCs proteome (Figure 1D).

Moreover, we identified five expression clusters based on the dynamic proteomic profile (Figure S1A and Table S1). A matrisome-enriched cluster, which rose on day 8 and included proteins related to ECM organization and collagen metabolism (Figure 1E and S1B). A metabolic cluster containing proteins, hereafter referred to as met-proteins, involved in vesicular processes and cell commitment which was up-regulated on days 4-8, (Figure 1F and S1B). A proliferation and pluripotency clusters, which contained proteins down-regulated during differentiation (Figure 1G-H, S1B). And a basal-proteins cluster, containing proteins that varied minimally involved in basal cellular processes (Figure 1I, S1B). Altogether, this broad characterization enables the construction of a proteomic roadmap of the differentiation of dHSCs from iPSCs.

### dHSCs resemble human primary HSCs proteome identity

We next evaluated the similarity between dHSCs and primary human HSCs (pHSCs) isolated from liver donors. We found that both populations shared more than 80% of their proteome profiles (Figure 1J) and by interrogating the proteomic data with published gene signatures of quiescent, activated, and stellate cell phenotypes (Zhang et al., 2016)^19^, we observed that during differentiation, dHSCs gained expression of pathways related to quiescence, activation, as well as pan-stellate cell phenotype (Figure 1K-M). Overall, these data confirms that iPSCs progressively acquire the stellate phenotype along differentiation, thus enabling the use of dHSCs to further investigate their role in cell commitment and disease.

### Cell trajectory analysis of dHSCs differentiation identifies RORA as a potential driver of HSCs differentiation

Principal component analysis (PCA) of the proteome differentiation profile showed that samples distribute along the differentiation process in a time-dependent manner (Figure 2A). Proteins comprised in PC1 explained stellate cell commitment across time, as cells gained in proteins related to collagen metabolism, ECM organization, or wound healing processes.

**Figure 2:**
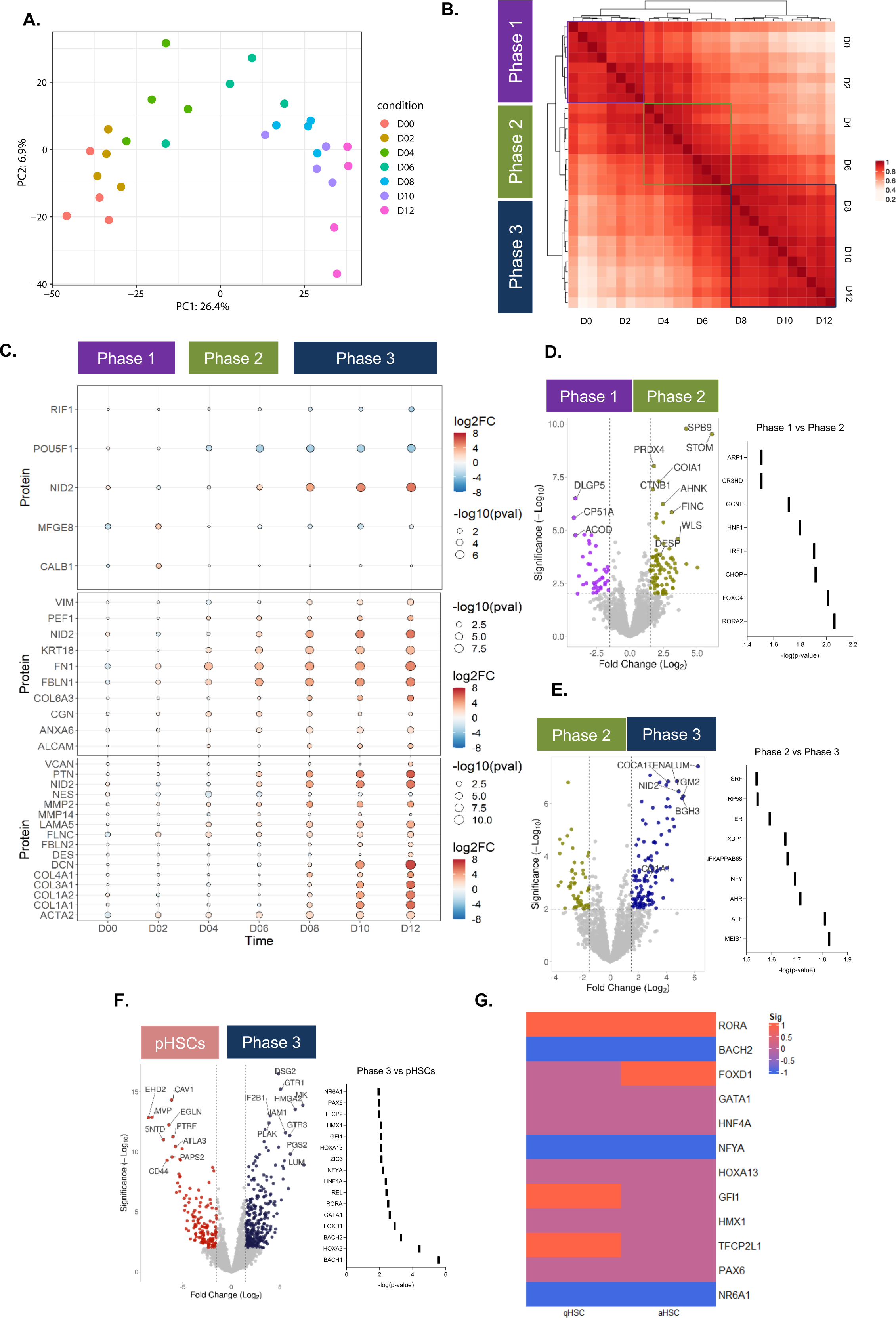
Cell trajectory analysis of dHSCs differentiation discovers three stages of maturation and identifies RORA as a potential driver of stellate cell differentiation. (A) Principal component analysis of the proteome time course showed that there is a partition of data in a time dependent manner. (B) Pearson correlation analysis revealed that differentiation occurs in three stages: Phase 1, from D0 (day 0) to D4 (day 4); phase 2 from D4 (day 4) to D8 (day 8) and final maturation phase (phase 3) from D8 (day 8) to D12 (day 12). (C) Dotplot showing the enrichment of proteins across differentiation: pluripotency and mesoderm proteins (POU5F1, RIF1, MFGE8 and CALB1) are enriched in phase 1 (from day 0 to day 4); in phase 2, mesenchymal and mesothelial proteins together with fetal HSCs started to be enriched (VIM, PEF1, KRT18, FN1, FBLN1, COL6A3, CGN, ANXA6 and ALCAM); finally, in phase 3 mature HSCs markers are enriched (VCAN, PTN, NES, MMP2, MMP14, LAMA5, FLNC, FBLN2, DES, DCN, COL4A1, COL3A1, COL1A2, COL1A1 and ACTA2). (D) Volcano plot of the comparison between phases 1 and 2 shows an enrichment in mesothelial markers (DESP, DSG2, PRDX4 and WLS) and in silico predicted TFs regulating this transition. (E) The comparison between phases 2 and 3 shows an increased in collagens and ECM proteins (COL5A1, LUM, TNC, TGFBI and NID2) and in silico predicted TFs modulating this transition. (F) Volcano plot showing differential protein expression between phase 3 (dHSCs) and primary HSCs and *in silico* predicted TFs modulating the adult stellate cell phenotype. (G) Transcriptomic data of primary activated and quiescent stellate cells in comparison to dHSCs of the predicted in silico TFs modulating the stellate adult phenotype.

To further dissect the progression of the differentiation, we performed Pearson’s correlation analysis showing that proteome data is divided into three phases (Figure 2B). Phase 1 included days 0 to 2 and was characterized by the expression of pluri- and prep-proteins, such as POU5F1 and RIF1 (Figure 2C) and early mesoderm proteins such as CALB1 and MFG-E8. In Phase 2, from day 4 to day 6, cells started to express met-proteins and mesenchymal (FN1, ANXA6, PEF1, and VIM), fetal liver mesothelial markers (FLNC, KRT18, CGN, and ALCAM), and fetal HSC markers such as COL6A3, NID2, FBLN, and PTN (Figure 2C). Finally, phase 3, from days 8 to day 12, was characterized by the final mature population of dHSCs, which express matrisome-proteins and mature HSCs markers, such as ACTA2, COL1A1, DES, and MMP2 (Figure 2C).

When comparing the expression profiles between phases, we observed that proteins differentially expressed from Phase 1 to Phase 2 were related to characteristics of mesenchymal and mesothelial cells such as desmosomes or tight junction formation (DESP, DSG2, PRDX4) (Figure 2D). Proteins differentially expressed between phases 2 and 3 were related to collagen and ECM organization (COL5A1, LUM, TNC, TGFBI, and NID2), which are typically expressed in HSCs, reinforcing the notion that mature HSCs appear in phase 3 (Figure 2E). These results indicate that the HSCs differentiation takes place in three phases mimicking key stages of embryonic development.

Next, we assessed which TF were involved in the trajectory of HSCs from cell differentiation to cell identity, starting from the TFs regulating the transition between the different phases of differentiation identified previously. We found that RORA was the most significant TF predicted to be regulating the transition between phases 1 and 2, indicating a possible role of RORA in mesoderm commitment (Figure 2D and Table S2A). Meis homeobox 1 (MEIS1) was predicted to regulate the transition from phases 2 to 3 (Figure 2E and Table S2B). Next, we assessed which TF were involved in the identity of the stellate cells by comparing Phase 3 cells with pHSCs (Figure 2F and Table S2C). Moreover, to ensure that the selected TF were relevant in primary human cells, we interrogated transcriptomic data from quiescent and activated primary human HSCs (GSE90525 & GSE67664). This analysis confirmed that RORA was the only TF up regulated in both quiescent and activated pHSCs when compared to dHSCs, thus suggesting its potential role in the acquisition the HSCs phenotype (fold change between dHSCs vs. qHSCs is 3.20; FC dHSCs vs. aHSCs is 1.60) (Figure 2G).

### RORA facilitates the dHSCs mature phenotype by improving mesoderm commitment

RORA is a member of the steroid/thyroid hormone receptor superfamily of TFs expressed in several tissues, including the liver, where is known to regulate genes involved in hepatocyte lipid, cholesterol and glucose metabolism. RORA plays a major role in cellular development and differentiation of multiple tissues^20,21^, but the role in HSCs physiology is unknown. During dHSCs differentiation, the gene expression of RORA showed a significant increase at day 4 of differentiation and progressively declined at days 6, 8, 10, lowering the pHSCs expression at day 12 (Figure 3A).

**Figure 3:**
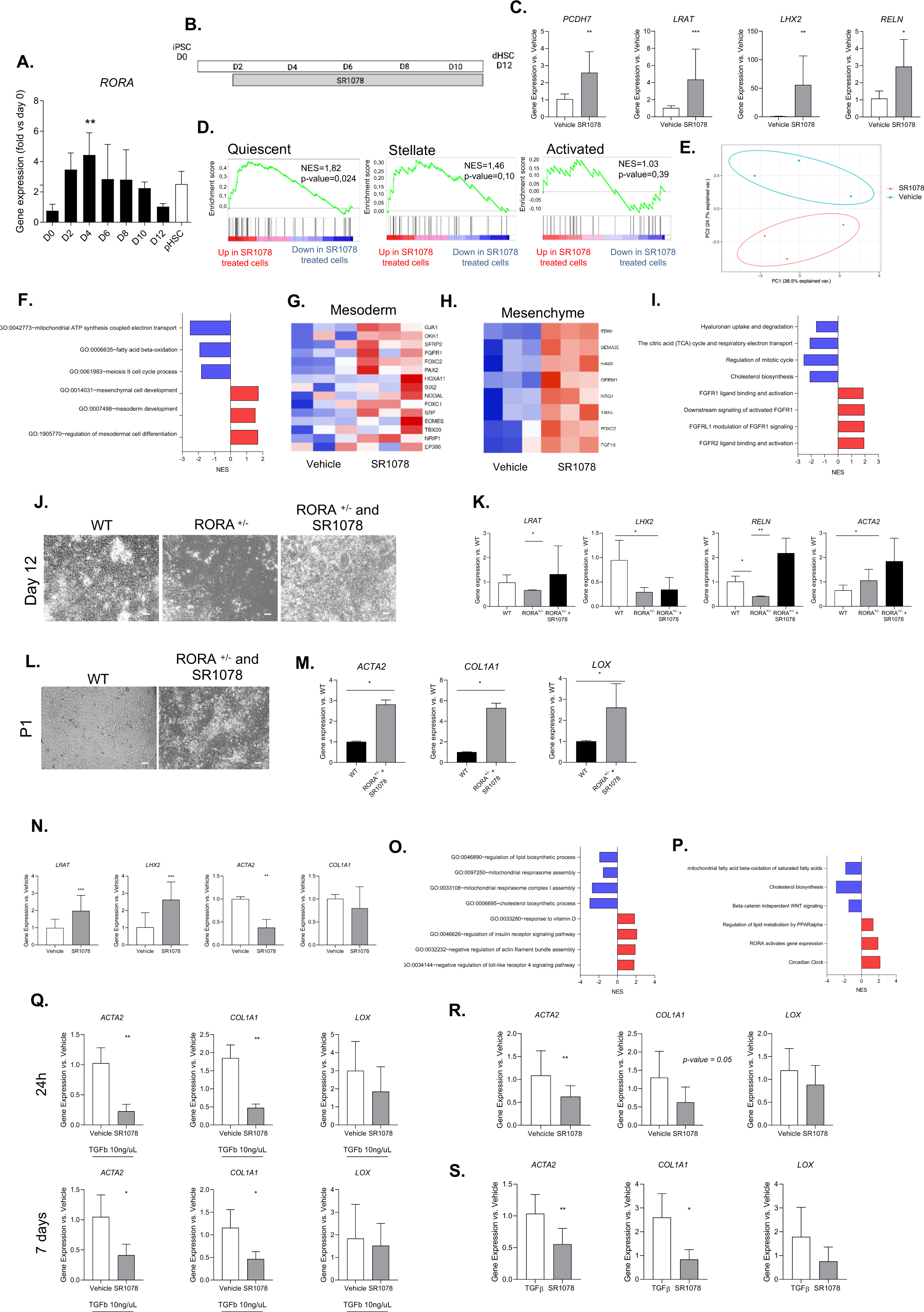
RAR-related orphan receptor A is a potential driver of the dHSCs phenotype by improving mesoderm commitment and modulating dHSCs activation state. (A) RORA gene expression across differentiation in comparison to pHSCs. The experiment was performed using three independent differentiations and three pHSCs. (B) Schematic illustration of the experimental design of SR1078 treatment across differentiation. Cells were treated at 0.1mM from day 2 onwards with the RORa agonist (SR1078). (C) Gene expression analysis of three independent differentiations treated with SR1078 in comparison to the vehicle group for stellate cell markers (*PCDH7, LRAT, LHX2 and RELN*). (D) GSEAs of reported gene signatures for quiescent, pan- and activated stellate cells. (E) PCA showing transcriptomic differences between cells treated with the RORA agonist from day 2 compared to the untreated group. (F) GOs upregulated (red) and downregulated (blue) in cells treated with the RORA agonist. (G) Heatmap of mesoderm markers in treated and untreated cells along differentiation with SR1078. (H) Heatmap of mesenchymal markers in treated and untreated cells along differentiation with SR1078. (I) Reactome pathways upregulated (red) and downregulated (blue) in cells treated with the RORA agonist. (J) Representative microscopy images of dHSCs from WT, iPSC-RORA^+/-^ and rescued iPSC-RORA^+/-^ with the RORA agonist at day 12. (K) Gene expression of stellate cell markers (*LRAT, LHX2, RELN* and *ACTA2*) of WT, iPSC-RORA^+/-^ and rescued iPSC-RORA^+/-^ with the RORA agonist at day 12. Scale bar represents 100μm. (L) Representative microscopy images of passaged (P1) dHSCs from WT and iPSC-RORA^+/-^ SR1078 treated. Scale bar represents 100μm. (M) Gene expression of activated stellate cell markers (*ACTA2, COL1A1 and LOX*). (N-O) Schematic illustration of the two in vitro activation models and the treatment with the RORA agonist. (P) Gene expression of quiescent (*LRAT* and *LHX2*) and activated (*ACTA2 and COL1A1*) markers in cells treated with the RORA agonist at passage. (Q) GOs upregulated (red) and downregulated (blue) in treated cells with the RORA agonist at passage. (R) Reactome pathways upregulated (red) and downregulated (blue) in cells treated with the RORA agonist at passage. (S) Gene expression of activated (markers in cells treated with the RORA agonist at passage, after TGFβ stimulation (10ng/μL) during 24h and 7 days. (T) Gene expression of activated markers (*ACTA2, COL1A1 and LOX*) in liver spheroids after SR1078 3mM treatment during 24h. (U) Gene expression of activated markers (*ACTA2, COL1A1 and LOX*) in liver spheroids after TGFb stimuli during 24h and SR1078 3mM treatment during 24h more. Significant differences are indicated as *p<0.05.

To assess the role of RORA across the differentiation, we induced RORA activity using the agonist SR1078 from day 2 until the end of the differentiation (day 12) (Figure 3B). While the RORA agonist did not change the morphology of dHSC (Figure S2A), we found that it increased the expression of mature HSC markers such as *PCDH7*, *LRAT*, *LHX2* and *RELN* (Figure 3C). At transcriptome level, the GSEA analysis of gene signatures of quiescent, activated and hepatic stellate phenotype (Zhang et al., 2016), revealed that RORA agonism promoted the quiescent gene signature (NES=1.82; p-value=0.024) (Figure 3D), while maintaining the ability to respond to TGFβ (Figure S2B). Moreover, as shown by differential clustering in the PCA (Figure 3E), RORA agonism induced transcriptomic changes in dHSCs, also reflected by the enrichment in biological processes and markers related to mesoderm (i.e.*EOMES*, *SRF* or *GJA1*) and mesenchyme development (i.e. *GREM1*, *HAS2* or *FGF8*) (Figure 3F-H) and reduced enrichment in the biological processes related to cell proliferation (Figure 3F and Table S3A). The functional transcriptomic analysis of differentially expressed genes in dHSCs treated with RORA agonist, identified enriched Reactome pathways such as FGFs signalling and a reduction of beta-oxidation, tricarboxylic acid cycle and mitotic cycle (Figure 3I and Table S3B), suggesting a role for RORA in regulating cell metabolism during dHSCs differentiation. Overall, these data indicate that RORA might act as a transcriptional regulator of the mesoderm and mesenchymal specification and dHSC differentiation.

### Functional role of RORA in mesoderm and mesenchymal specification

To further dissect the role of RORA in dHSCs differentiation, we generated three clones of heterozygous RORA-knockout iPSCs (RORA^+/-^) by CRISPR-Cas9 mimicking the mutation presented in staggerer mice (sg/sg), spontaneous KO mice for the RORA gene^22^. Generation of iPSCs RORA^+/-^ was confirmed by Sanger sequencing (Figure S2C). Cells showed a reduction in protein expression of RORA preserving pluripotency markers such as *SOX2*, *OCT4* and *NANOG* at both gene and protein expression (Figure S2D-F). Moreover, all the clones generated presented a normal karyotype, thus confirming the genome stability of the newly generated cell lines (Figure S2G). Interestingly, during the differentiation of RORA^+/-^ iPSCs we observed in all the clones tested with nearly 80% of the cells dying at day 4 (only results from clone 13 are shown) (Figure 3J). Survival cells that finalized the differentiation presented a lower expression of mature HSCs markers (*LHX2*, *LRAT*, *RELN* and *LRAT*) and an increase of activation genes such as *ACTA2* (Figure 3K). To rescue the cell mortality observed in RORA^+/-^ iPSCs, we treated the cells with the RORA agonist (SR1078) from day 2 and during the differentiation. The treatment rescued the mortality of RORA^+/-^ differentiating cells and the phenotype of dHSCs (Figure 3J-K). These results were confirmed by using a chemical antagonist of RORA (SR1001) in the differentiation of wild type (WT) iPSCs. SR1001 treated cells showed a spindle morphology with increased ECM production at the end of differentiation (Figure S2H-J), while reducing gene expression of the HSCs genes *RELN* and *LHX2* and increasing *COL1A1* and *ACTA2* (Figure S2J), together with increased collagen deposition, as assessed by Sirius Red staining (Figure S2I). Overall, these results position RORA as a transcriptional regulator of mesoderm and mesenchymal specification and key for the acquisition of the quiescent phenotype of dHSCs.

### RORA promotes a HSC quiescent phenotype and prevent cell activation

As the RORA agonist boosted the quiescent phenotype of dHSCs, and cells derived from iPSC-RORA^+/-^ cells had a higher activation profile (Figure 3L-M), we evaluated the effect of RORA on dHSCs activation and fibrosis using two *in vitro* models of dHSCs activation, one based on passage and the other based on TGFβ stimuli. Previously we have shown that passaged dHSCs are a good model to induce cell activation promoting a fibrogenic response (Figure S2K)^14^. dHSCs derived from WT iPSCs treated with the RORA agonist SR1078 for 24h, prevented activation of passaged dHSCs and decreased the expression of activation markers *ACTA2* and *COL1A1* while increased the expression of quiescent markers *LHX2* and *LRAT* (Figure 3N). On the contrary, cells treated with the RORA antagonist (SR1001) after passage, increased their activation profile (Figure S2L-M). Transcriptomic analysis of the passaged dHSCs with SR1078 treatment showed a reduction in GOs related to *de novo* lipogenesis and mitochondrial beta-oxidation of fatty acids (Figure 3O and Table S3C). In addition, as shown in Figure 3P and Table S3D, Reactome analysis showed changes in metabolic pathways, thereby suggesting a remodeling of the metabolic state in the dHSCs (Figure 3R).

Next, we evaluated the effect of RORA on dHSCs response to TGFβ, as a second activation model^17^. dHSCs were treated with TGFβ for 24h or 7 days while the SR1078 RORA agonist was added for the last 24h. dHSCs treated with SR1078 reduced both short and long-term TGFβ-activation by decreasing *ACTA2*, *COL1A1* and *LOX* (Figure 3Q). To study the effect of RORA in a complex *in vitro* system, we evaluated the effect of SR1078 in 3D liver spheroids of HepG2 and dHSCs. Incubation of spheroids with SR1078 for 24h reduced the expression of *COL1A1*, *ACTA2* and *LOX* (Figure 3R). Moreover, liver spheroids incubated with TGFβ and treated with SR1078 also showed a reduced expression of activation markers (*COL1A1* and *ACTA2*) (Figure 3S) as well. Overall, these results suggest that RORA plays a role in the deactivation of dHSCs and the maintenance of the quiescent dHSCs phenotype, which can be mediated by blocking metabolic adaptations required for HSCs activation.

### RORA regulates cell metabolism during mesoderm differentiation and dHSCs activation

Since our data showed that RORA may be responsible for the modulation of the metabolic state of the cells along dHSCs differentiation and activation, we evaluated dHSCs metabolic reprogramming in the presence and absence of RORA, both during differentiation and activation. Previous reports showed that cells exiting pluripotency and differentiating to mesoderm undergo a metabolic switch increasing OXPHOS and reducing glycolytic flux^23^. Similarly, activation of HSCs is dependent upon induction of glycolysis^10^, glutaminolysis^11^ and *de novo* lipogenesis^12^. With the use of real-time Seahorse cell metabolic analysis, we found that cells from iPSC-RORA^+/-^ differentiation have increased mitochondrial respiration at day 4 (basal, ATP-coupled and maximal respiration) (Figure 4A and B). In addition, adenosine triphosphate (ATP) production from glycolysis was also increased in these cells, (Figure 4A-C), in parallel with to an increase in intracellular glucose levels (Figure 4D), thus maintaining the metabolic profile of undifferentiated cells. Also, iPSC-RORA^+/-^ cells expressed higher levels of genes related to glycolytic flux (*ALDOB* and *GS6P*) and a decreased expression of *ACACA* (Figure 4E). Moreover, the expression of mesoderm (EOMES) and mesenchymal (VIM) markers was reduced at both protein and gene expression in iPSC-RORA^+/-^ cells at day 4 of the differentiation as well (Figure 4E-F) when compared to WT iPSCs. However, when iPSC-RORA^+/-^ cells were treated with RORA agonist, they showed a reduction in intracellular glucose and glycolysis-dependent ATP production (Figure 4C-D), glycolytic gene expression (Figure 4E) and glucose consumption from the media (Figure 4D), thus confirming that iPSC-RORA^+/-^ cells are not able to initiate mesoderm commitment and retain the metabolic profile of undifferentiated cells.

**Figure 4:**
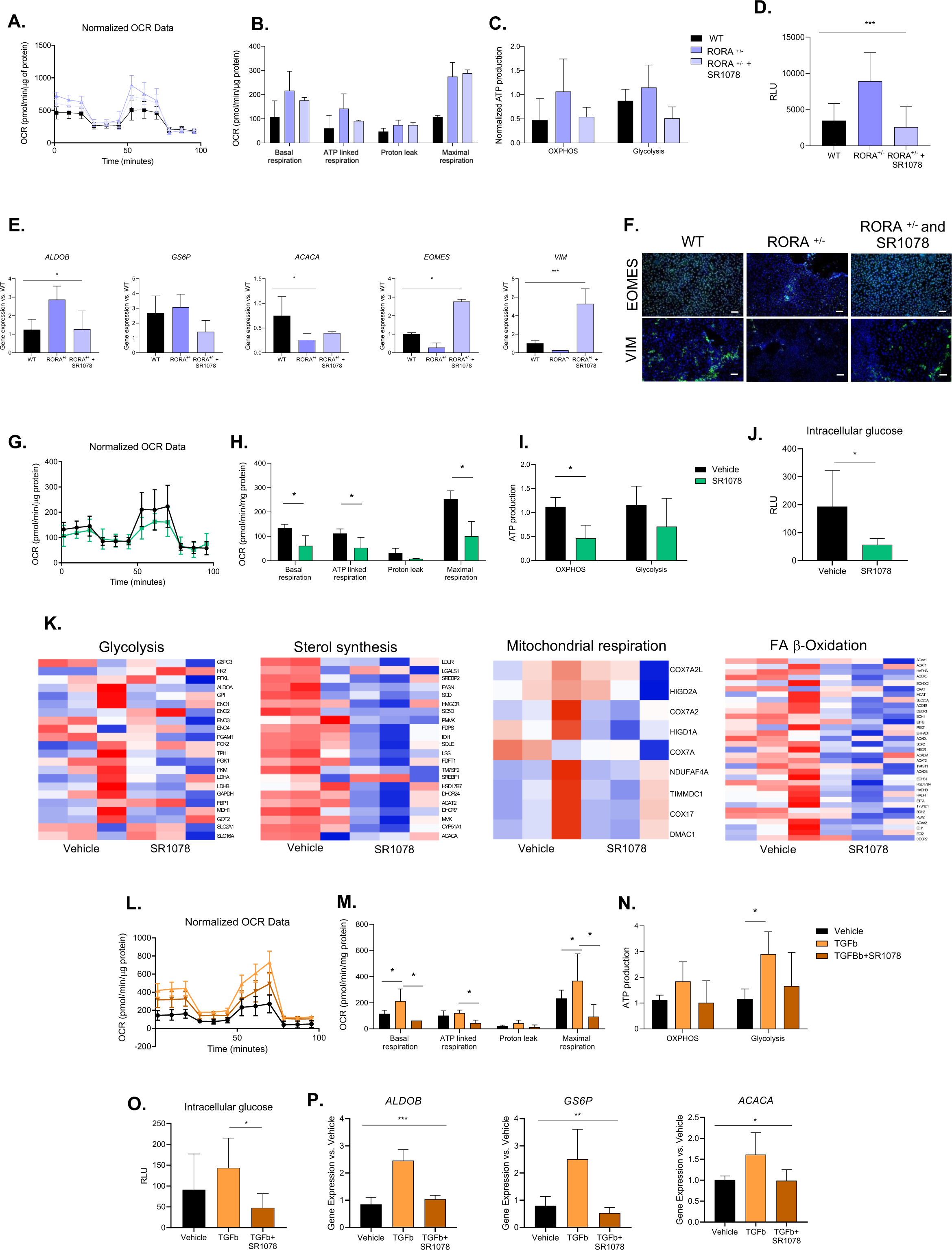
RORA regulates cell metabolism during mesodermal differentiation and dHSCs activation. (A) Oxygen consumption rate (OCR) on day 4 WT iPSC and iPSC RORA^+/-^ treated and untreated from day 2. (B) OCR parameters of day 4 iPSC WT and iPSC RORA^+/-^ treated and untreated from day 2. (C) Normalized ATP production from OXPHOS and glycolysis of day 4 WT iPSC and iPSC RORA^+/-^ treated and untreated from day 2. (D) Glucose consumption at day 4 of WT iPSC and iPSC RORA^+/-^ differentiations treated and untreated from day 2. (E) Gene expression of key glycolytic and lipid synthesis markers (*ALDOB*, *GS6P*, *ACACA*) and mesoderm and mesenchymal markers (*EOMES* and *VIM*). (F) Immunofluorescence of mesoderm (EOMES) and mesenchymal (VIM) on day 4 of differentiated cells from WT, RORA^+/-^ and RORA^+/-^ treated with the RORA agonist from day 2. Scale bars represent 100μm. (G) OCR passaged dHSCs treated with the RORA agonist for 24 hours. (H) OCR parameters of passaged dHSCs treated with the RORA agonist for 24 hours. (I) Normalized ATP production from OXPHOS and glycolysis of passaged cells treated and untreated with RORA agonist for 24 hours. (J) Glucose consumption at passaged cells treated and untreated with RORA agonist for 24 hours. (K) Heatmap of glycolytic, sterol synthesis, mitochondrial respiration and fatty acid beta-oxidation of passaged cells treated and untreated cells with RORA agonist for 24 hours. (L) OCR of passaged cells treated with TGFβ 10ng/mL for 24 hours and rescued with RORA agonist treatment for 24 hours more. (M) OCR parameters of passaged cells treated with TGFβ 10ng/mL for 24 hours and rescued with RORA agonist treatment for 24 hours more. (N) Normalized ATP production from OXPHOS and glycolysis of TGFβ activation model rescued with the RORA agonist for 24 hours. (O) Intracellular glucose levels of TGFβ activation model rescued with the RORA agonist for 24 hours. (P) Gene expression of key glycolytic and lipid synthesis markers (*ALDOB*, *GS6P* and *ACACA*) of the TGFβ activation model rescued with the RORA agonist for 24 hours. Significant differences are indicated as *p<0.05.

Similarly, treatment with the RORA agonist in both activation models previously used reduced mitochondrial respiration, OXPHOS-dependent ATP production, glycolysis-dependent ATP production as well as intracellular glucose (Figure 4G-J, and 4L-O). Moreover, genes associated with glycolysis and lipid synthesis that were increased with activation, were found to be reduced after treatment with SR1078 (Figure 4K and 4P). Altogether, these results indicate that RORA acts as a metabolic modulator in dHSCs trajectory which impacts the phenotype of dHSCs at different stages, including initial cell commitment and the quiescent cell identity at the end of differentiation.

### RORA is involved in fibrogenic response *in vivo*

To evaluate the role of RORA in HSCs activation and liver fibrosis, we used a RORA-deficient strain, known as staggerer mice, hereafter sg/sg. We found no significant differences in baseline fibrosis of wildtype (WT) and sg/sg livers as shown by the hydroxyproline assay, and we detected a very slight increase in the expression of *Acta2 and Timp1* activation markers (Figure S3A and B).

Next, we evaluated the fibrogenic response of the livers of sg/sg mice by inducing liver fibrosis using CCl_4_ for 4 weeks (Figure 5A). The livers of the sg/sg group presented an increased fibrotic content (Sirius Red) coupled with an increased in alpha smooth muscle actin (αSMA) staining as compared to their littermate WT counterparts (Figure 5B). In agreement with previous reports showing increased hepatocyte injury in RORA-KO mice fed with a high-fat diet^24^, by hematoxylin and Eosin (H&E) staining we observed an increased tissue damage in the pericentral hepatocytes (Figure 5B) in the sg/sg group. The levels of alanine transaminase (ALT), aspartate transaminase (AST), phosphatase alkaline (AP) and lactate dehydrogenase (LDH) in sg/sg mice were also higher than in the WT counterparts (Figure S3C). Moreover, gene expression of fibrogenic markers such as *Acta2*, *Col1a1*, *Col1a2*, *Timp1* and *Fn1* were significantly increased in the mutant group (Figure 5C), with an increased neutrophil content (Figure S3D), and no significant differences in macrophages (F4/80) or lymphocytes (CD3) (Figure S3D). No differences in proliferation were observed between groups (Figure S3D). Altogether, these results reproduce the *in vitro* models confirming the role of RORA in fibrogenic response.

**Figure 5:**
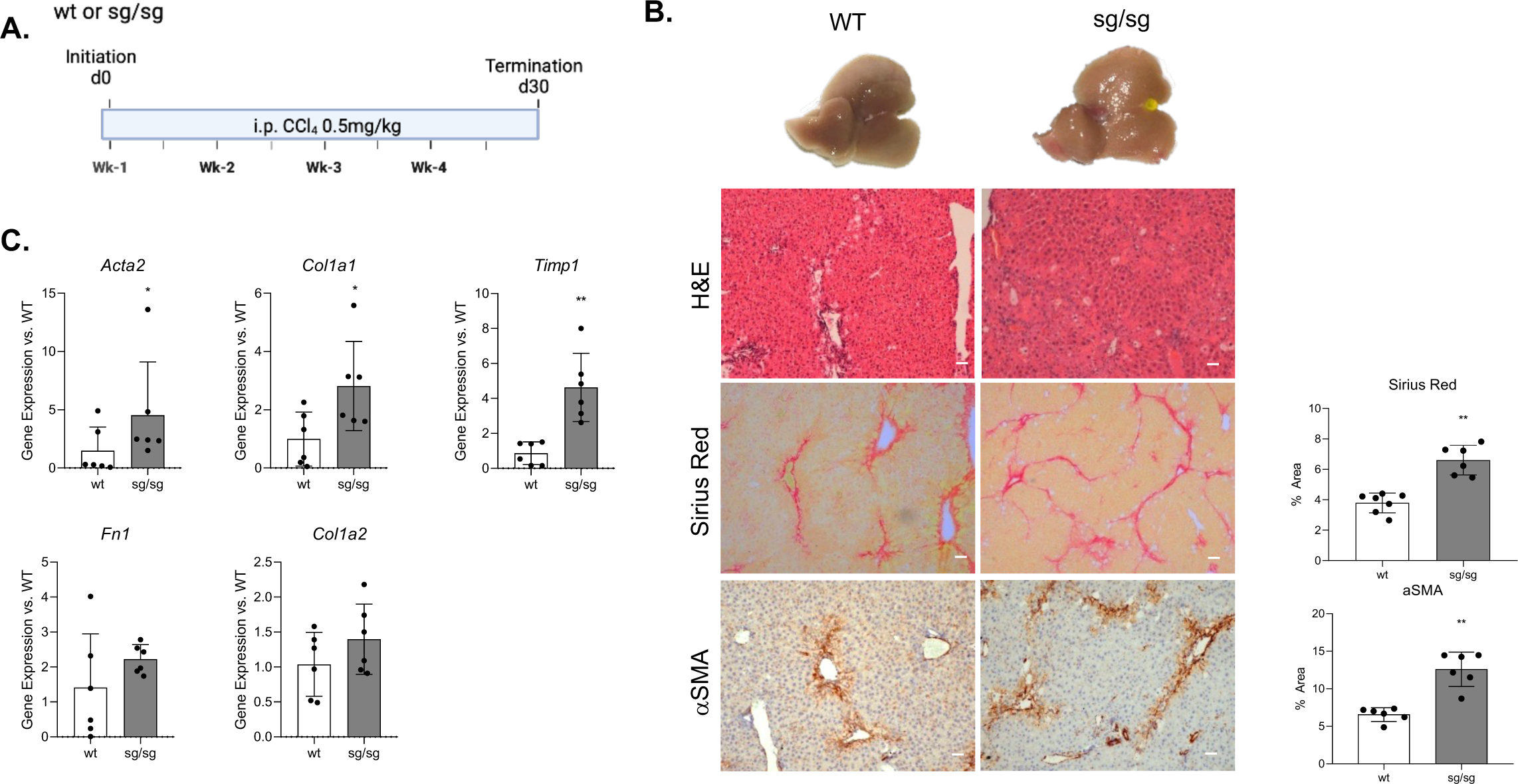
RORA controls the activation of HSCs and contributes to the development of liver fibrosis. (A) Schematic overview of a CCl_4_ fibrotic liver model in staggerer mice. (B) Representative images of the livers after 4 weeks of CCl_4_ in staggerer and wildtype, H&E staining and Sirius Red staining for the staggerer and wildtype counterparts and immunohistochemistry of αSMA. Scale bars represent 100μm. (C) Gene expression of fibrogenic and HSCs activation markers (*Acta2, Col1a1, Timp1, Fn1* and *Col1a2*) in 7 staggerer and 7 WT mice after CCl_4_ treatment. Significant differences are indicated as *p<0.05.

### The RORA agonist reduces liver fibrosis in multiorgan fibrosis models

To explore the translational potential of targeting RORA in fibrogenic diseases, we administered the RORA agonist SR1078 to CCl4-treated mice (Figure 6A). Mice treated with SR1078 increased *Rora* gene expression (Figure 6B). H&E staining showed reduced tissue damage coupled with reduced collagen deposition and αSMA staining (Figure 6C). Likewise, we observed a reduced gene expression of fibrogenic and HSCs activation markers such as *Acta2*, *Col1a1*, *Timp1*, *Fn1*, *Col1a2*, *Pparg*, *Mmp2*, *Mmp9*, *Mmp12* and *Timp1* (Figure 6D and Figure S4A). Immunohistochemistry for F4/80, MPO and CD3 showed a non-significant reduction of inflammatory cell populations in mice treated with SR1078 (Figure S4C). In accordance, gene expression for *Il1b*, *Tnfa* and *Il6* inflammatory cytokines presented a non-significant reduction (Figure S4A). Moreover, parenchyma liver damage was reduced in SR1078 treated mice, showing a reduction of ALT, AST and LDH serological levels (Figure S4B). Also, it reduced the proliferative activity of liver parenchyma as shown by Ki67 staining (Figure S4C). These results suggest that the RORA agonist has a dual effect in mitigating liver damage and reducing HSCs activation and fibrosis.

**Figure 6:**
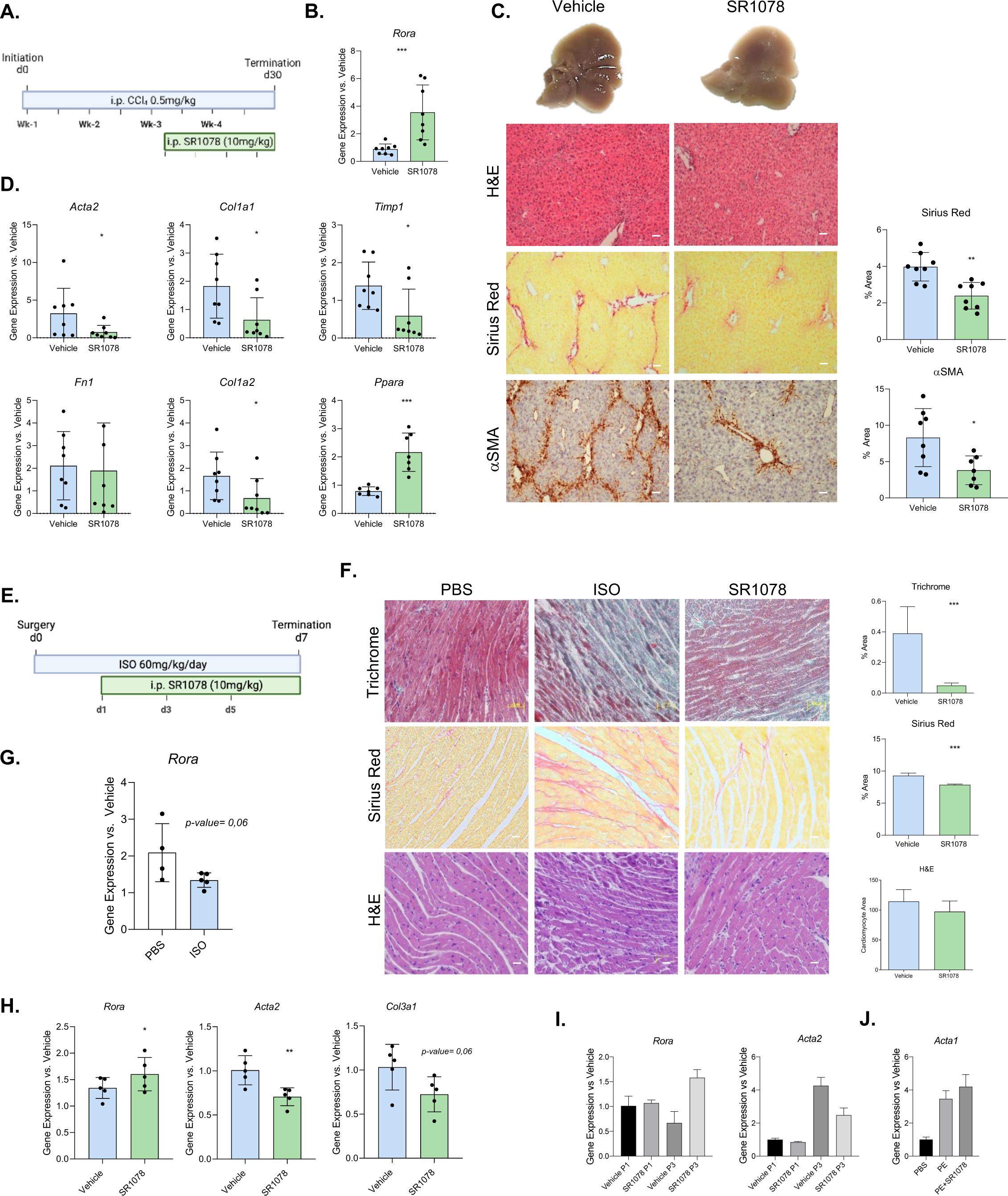
dHSCs trajectory analysis is a reliable technique for finding anti-fibrogenic targets. (A) Illustration of the experimental design of a CCl_4_ model treated with SR1078. (B) RORA gene expression of CCl_4_ model treated with SR1078. (C) Representative images of the livers after CCl_4_ treatment and SR1078, H&E staining and Sirius Red staining for the untreated (Vehicle) and treated (SR1078) group and immunohistochemistry of αSMA. Scale bars represent 100μm. (D) Gene expression of fibrogenic and HSCs activation markers in 7 untreated (Vehicle) and 7 treated (SR1078) mice (*Acta2, Col1a1, Col1a2, Timp1* and *Fn1*). Significant differences are indicated as *p<0.05. (E) Illustration of the experimental design of Isoproterenol cardiac injury model. (F) Representative images of the hearts of vehicle, isoproterenol (ISO) and ISO and RORA agonist group of Masson’s trichrome, Sirius Red and H&E staining. Scale bars represent 100μm.(G) *Rora* gene expression of vehicle and ISO group. (H) Gene expression of *Rora* and fibrogenic markers (*Acta2* and *Col3a1*) and the hypertrophy marker *Acta1* in ISO and ISO and RORA agonist group. (I) Gene expression of Rora, activation markers (Acta2) and cardiac hypertrophy markers (Nppa and Pdk4) in in vitro culture of cardiac fibroblast and cardiomyocytes. * Significant differences are indicated as *p<0.05.

HSCs are liver pericytes showing important phenotypic and functional similarities with pericytes from other organs and participating in wound healing response. Hence, we evaluated the effect of RORA agonism in extrahepatic wound healing response models. First, we tested the effect of the SR1078 RORA agonist in an isoproterenol-induced cardiac hypertrophy mice model (Figure 6E). The model showed an increased in cardiac fibrosis (Figure 6F) and reduction in the cardiac expression of *Rora* (Figure 6G). Treatment with the RORA agonist promoted the expression of *Rora* and reduced cardiac fibrogenic genes (*Col3a1* and *Acta2*) (Figure 6H) and ECM deposition as assessed by Masson’s trichrome and Sirius Red (Figure 6F). Echography analysis showed no differences in cardiac structure and function between the vehicle and the treated group (Table 1). Moreover, RORA treatment did not alter cardiomyocyte hypertrophy as shown by no differences in either the expression levels of the hypertrophy marker *Acta1* nor in the cardiomyocyte area (Figure 6F), suggesting a positive effect of RORA agonism against cardiac fibrosis independently of hypertrophy development. The antifibrogenic role of RORA was confirmed using primary culture of neonatal cardiomyocytes (NCMs) and fibroblasts (NCFs). Cultured fibroblasts to passage 3 reduced *Rora* expression while increasing *Acta2,* while RORA treatment reduced this expression, confirming its anti-fibrogenic effect (Figure 6I). By contrast, in NCMs, the RORA agonist did not alter cardiomyocyte area or the expression levels of the *Acta1* hypertrophy marker in the presence of the hypertrophic agent phenylephrine (PE) confirming the absence of anti-hypertrophic effects *in vitro* (Figure 6J). Second, we explored the antifibrogenic potential of RORA in a kidney damage model based on a unilateral ureteral obstruction (UUO) intervention (Figure S4D). The UUO model reduced the expression of *Rora* and promoted kidney fibrosis (Figure S4E). The RORA agonist-treated group showed a reduction in hydroxyproline content, indicating a reduced collagen deposition (Figure S4F), but a non-significant reduction of αSMA expression (Figure S4G). Finally, treated mice did not showed a non-significant reduction in the gene expression of *Acta2* and *Col1a1* and an increased expression of *Rora* (Figure S4H). These results indicate that pharmacological intervention boosting RORA activity is a potential strategy for multiorgan fibrogenesis mitigation.

**Table 1:**
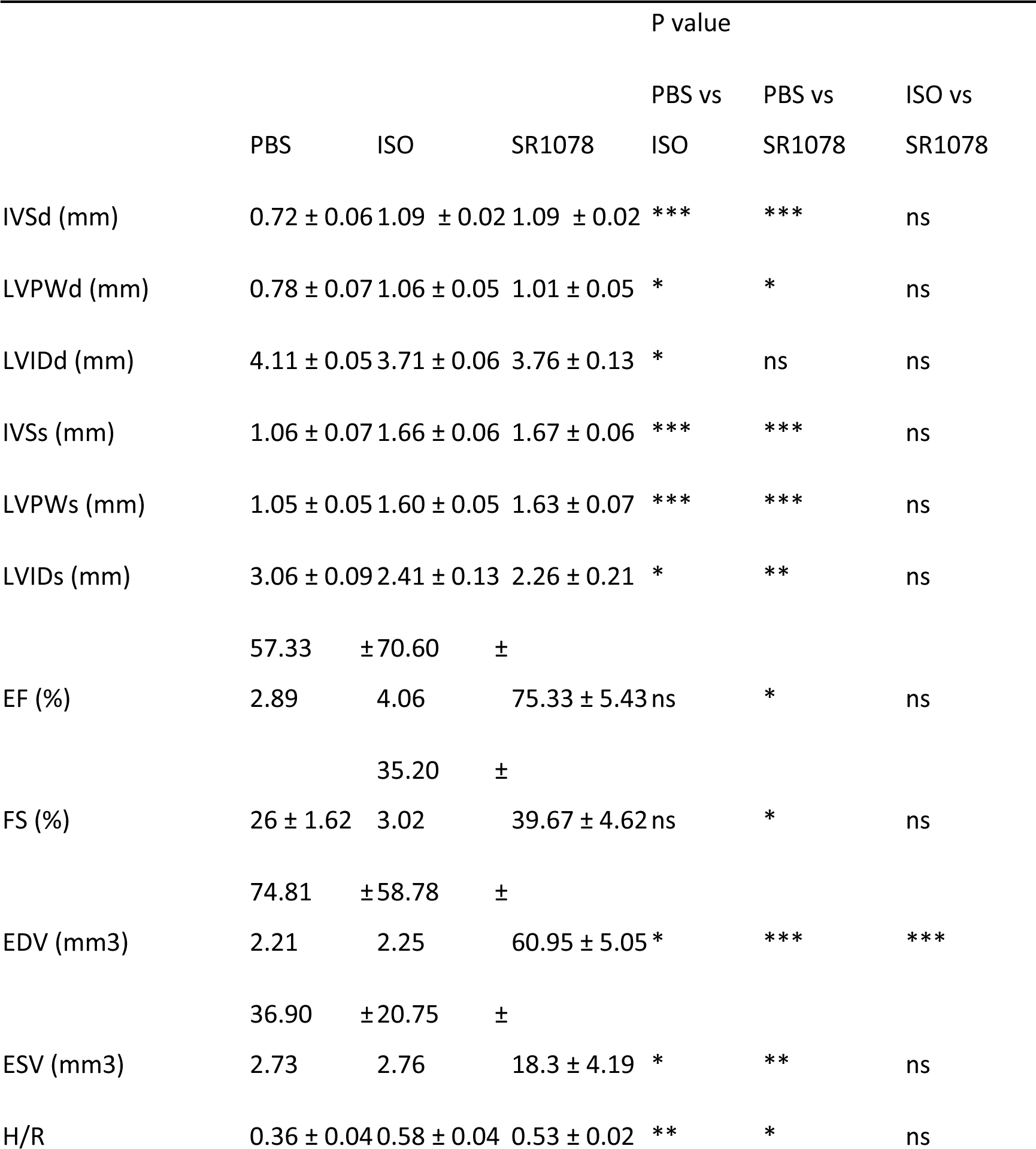
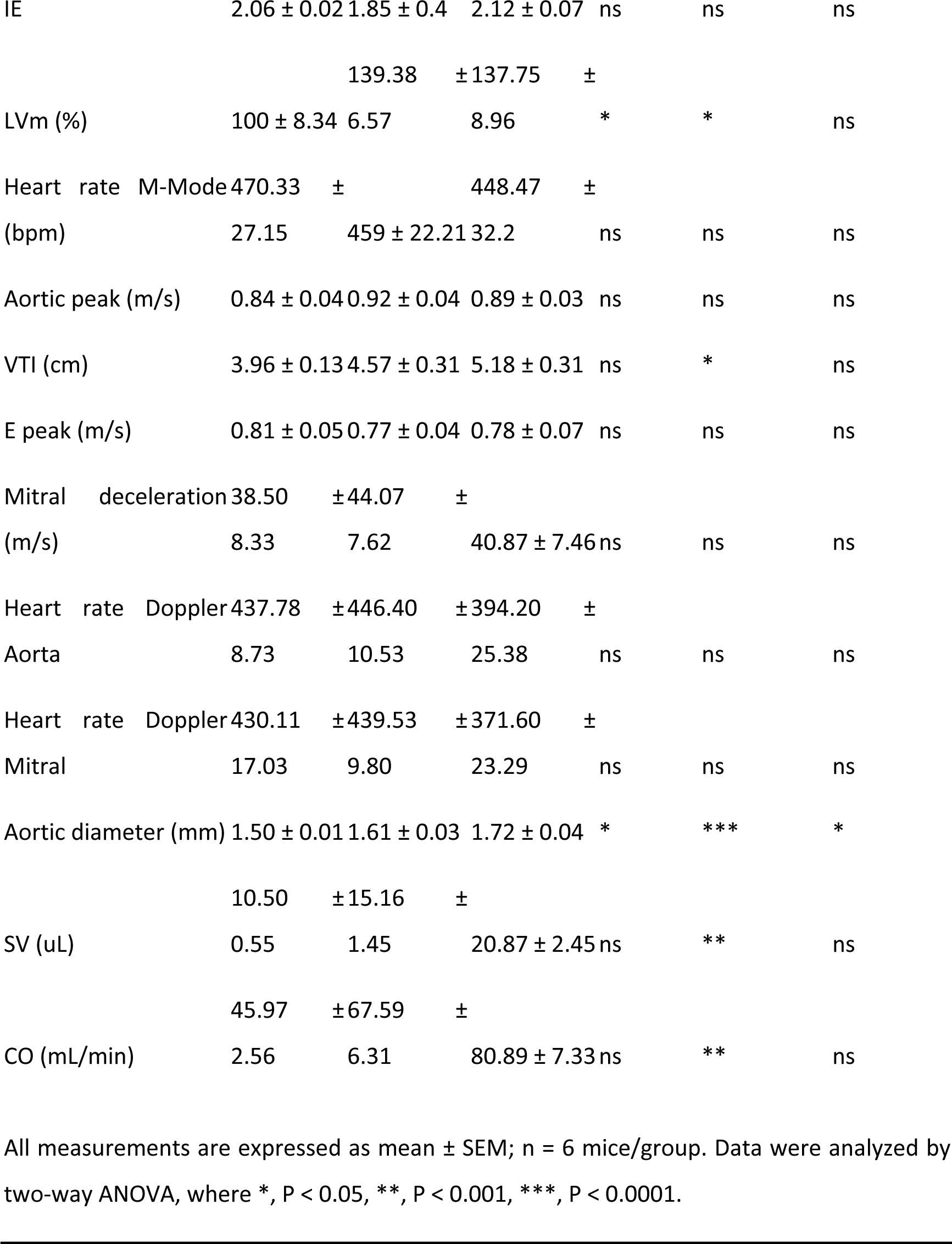
Echocardiographic data from WT animals treated with the RORA agonist (SR1078) after isoproterenol-induced hypertrophy. LVIDd, Left ventricular internal diameter during diastole; LVIDs, Left ventricular internal diameter during systole; IVSd, internal ventricular septum diameter during diastole; IVSs, internal ventricular septum diameter during systole; LVPWd, Left ventricular posterior wall thickness diameter during diastole; LVPWs, Left ventricular posterior wall thickness diameter during systole. EDV, end-diastolic volume; ESV, end-systolic volume; EF, ejection fraction; FS, fractional shortening; H/R, ventricular thickness/radius; IE, sphericity index; LVm, left ventricular mass; VTI, Velocity time integral; SV, stroke volume; CO, cardiac output.

### RORA gene expression correlates negatively with human liver fibrosis

Finally, to evaluate the relevance of our results in clinical settings, we evaluated RORA expression in cirrhotic patients, and we found that RORA protein and gene expression were both reduced in liver tissue from cirrhotic patients compared to controls at both gene and protein expression (Figure 7A and B). In addition, HSCs isolated from cirrhotic patients showed lower RORA expression than quiescent HSCs (FC=-1,88, p-value=0,02) (Figure 7C), according to our data showing RORA loss during HSCs activation. Next, we evaluated the correlation of RORA gene expression with fibrosis in a cohort of patients with different stages of chronic liver disease^25^. RORA gene expression was decreased in patients with Metavir F1-4 (Figure 7D) and correlated with the FIB4 fibrosis score (Figure 7E). Moreover, *RORA* expression negatively correlated with the expression of HSC activation and fibrogenic markers such as *ACTA2*, *COL1A1, LOXL1*, *TIMP1*, *VEGFC* and *TAGLN* (Figure 7F). Together with experimental *in vivo* and *in vitro* evidence, our findings support a role for RORA in pHSCs during the fibrogenic response, positioning RORA as a druggable target in chronic liver disorders.

**Figure 7:**
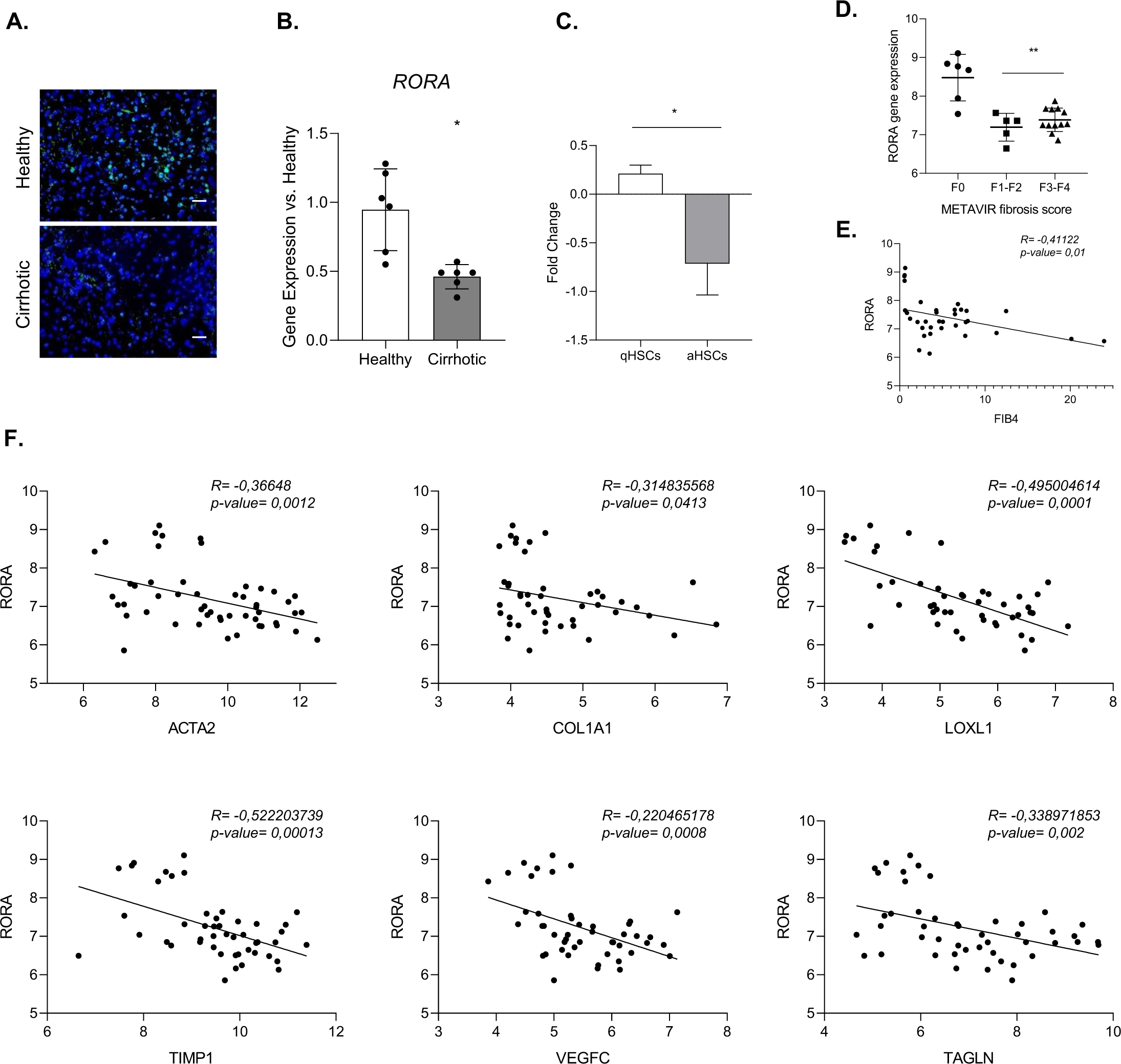
RORA is downregulated in a cohort of patients with liver disease and is negatively correlated with liver fibrosis. (A) Protein expression of RORA in cirrhotic patients is reduced in comparison to healthy group. Scale bars represent 100μm. (B) Gene expression of RORA in cirrhotic patients is reduced in comparison to healthy group. Six healthy livers and six cirrhotic livers were analyzed by quantitative polymerase chain reaction (qPCR). (C) Expression of RORA is reduced in primary activated HSCs in comparison to quiescent HSCs (GSE90525 & GSE67664). (D) RORA expression correlates negatively with METAVIR fibrosis score and FIB4 in a cohort of liver disease patients. (E) RORA correlates negatively with HSCs activation markers and fibrosis markers (*ACTA2, COL1A1, LOXL1, COL1A1, TIMP1, VEGFC* and *TAGLN*) in a cohort of liver disease patients. Significant differences are indicated as *p<0.05.

## Discussion

In this study we sought to improve our understanding of the trajectory of HSCs from development to disease and identify molecular drivers of cell identity and pathogenesis. Understanding HSCs paths in differentiation and activation is essential to delineate how cells acquire their phenotype and how this is lost during disease, thereby providing pathophysiological insights for the development of novel anti-fibrogenic treatments. Although cell trajectory upon disease can be studied using animal models, the translation to human samples is sometimes complex, and differentiation trajectories during embryo development are difficult to study. The use of differentiation protocols from human iPSCs towards adult cells together with relevant disease models are useful tools in this context, as we can study the whole spectrum of a cell, during differentiation, physiology and disease.

In this article, we performed a time resolving proteomic characterization of the differentiation of dHSCs, identifying a protein signature of dHSCs differentiation. The comprehensive proteomic data generated in this study will be a useful resource for researchers interested in understanding different aspects of the biology of human HSCs and fibrosis.

The differentiation process involves a major molecular switch from pluripotency to differentiated cells. Differentiation of iPSCs into dHSCs is associated with opposing albeit coordinated changes, silencing protein expression involved in cell proliferation and pluripotency, while acquiring the expression of proteins characteristic of the HSC phenotype and function. During dHSCs differentiation we detected stage specific markers reportedly expressed at different phases of the embryonic development of HSCs. In this regard, we show that differentiation occurs in three maturation stages, which express markers of mesoderm, mesothelial and immature HSCs. In this respect, the trajectory of dHSCs recapitulate the expression of protein markers described in the embryonic development of HSCs such as ALCAM, ANXA6, CGN, NES or DES ^3^. Unfortunately, we lacked a detailed understanding of the human development of HSCs, and therefore we cannot ensure that dHSCs differentiation is faithfully recapitulating all the features and the natural path of the human embryonic development.

One of the reasons behind the poor maturation of iPSC-derived cells is the divergence between primary cell development and iPSC-differentiation, which frequently deviate from the natural development path after embryonic stages, thus preventing the maturation of cells ^26^. In our study, we identified the role of RORA as a key transcriptional regulator involved in the mesoderm and mesenchymal specification of the cells and in the maintenance of their quiescent phenotype. Moreover, while deletion of RORA impairs early-stage differentiation, the activation of RORA improved dHSCs phenotype. Therefore, our study is in agreement with the notion that improving the intermediate specification steps of differentiation protocols may be a useful strategy to generate more mature adult cells. The study of the trajectory of dHSCs also highlighted the expression during differentiation of TF known to regulate HSCs biology and activation such as MEIS1, the role of which is not known in HSC development but is described to be involved in the activation of HSCs ^27^.

The trajectory analysis of dHSCs differentiation identified RORA not only as a regulator of mesoderm differentiation but also as a regulator of dHSCs identity and quiescence. The dHSCs derived from iPSC-RORA^+/-^, showed an increased activation phenotype. Also, pharmacological intervention in dHSCs and in a mouse model of liver fibrosis and extrahepatic fibrotic models show that activation of RORA prevents pericyte activation and fibrosis. In agreement with these results, human data from a cohort of chronic liver disease patients^28^ showed that the reduction of RORA is associated with the progression of fibrosis. Altogether, these results indicate that RORA has a dual role in HSC physiology, playing a role in HSCs differentiation and maturation but also preventing their activation, thus suppressing its fibrogenic response. Few reports have addressed the role of TF regulating HSCs activation during development. Both *Tcf21* and *Lhx2* are TF expressed during embryonic development of the liver, and their expression in HSCs is downregulated in activation and fibrosis ^13,1415^. Although their role at the different stages of HSCs maturation is not totally understood, these observations are consistent with our findings, suggesting that some TFs may play a role throughout development and disease. This highlights the potential of iPSC-differentiation approaches for investigating cell trajectories and they might be a powerful tool for understanding molecular drivers of differentiation and disease.

RORA is a pleiotropic TF involved in different biological functions and is also important for the correct differentiation of multiple cell types^29^. Fibrosis is an evolutionarily conserved mechanism developed by an organism to respond to chronic injury. Excessive fibrosis, however, leads to disruption of organ function and is a common feature of many chronic diseases, independently of the organ^30^. Thus, regardless of the organ, fibrosis shares many similar features, suggesting the existence of common core pathways^30^. We believe that dHSCs can be used as an archetype for pericytes, also in other tissues, as our results obtained in fibrosis models of extrahepatic organs indicate that RORA regulates not only HSCs activation but also other tissue pericytes and their role in fibrosis thereby suggesting that RORA could be an interesting target for the development of antifibrogenic treatments in different diseases and territories.

Metabolism reprogramming is key for correct embryonic development or the cell adaptation to stress or injury as it provides to each cell with the energetic requirements at the exact moment and is key for the correct embryonic development or the cell adaptation to stress or injury. For example, metabolic switching during mesoderm differentiation impairs the glycolytic flux towards a higher OXPHOS^31^. Similarly, activated myofibroblasts, including HSC in the liver, have been described to require a higher energetic demand during activation, because of increased proliferation, secretion of ECM, proteases and cytokines^32^. Transcriptomic data of activated dHSCs treated with the RORA agonist showed that there is a metabolic remodeling of the cells upon treatment, as they decreased beta-oxidation of lipids and glycolytic flux are decreased. For instance, fatty acid beta-oxidation is an important energy source for HSCs undergoing activation, as inhibition of mitochondrial fatty acid catabolism blocks HSC activation^33^. Moreover, acetyl-CoA carboxylase (ACC), a regulator of fatty acid beta-oxidation and *de novo* lipogenesis, has been implicated in HSC metabolic reprogramming during activation ^12^. In cells treated with the RORA agonist, ACC gene expression in dHSCs decreases. Also, activated dHSCs using two *in vitro* models, showed an increase in OXPHOS and glycolytic flux that was downregulated when treated with RORA, thus reducing the metabolic requirements of the dHSCs and preventing their activation. In addition, our results showed similar results at early stages of differentiation, as iPSC-RORA^+/-^ presented a metabolic profile characteristic of undifferentiated cells that was modified when treated with the RORA agonist, indicating that RORA regulates the metabolic activity during early stages of differentiation, promoting mesoderm differentiation. These results indicate that metabolic plasticity is required at different stages of the HSC trajectory and impairs the final phenotype of the cell. This metabolic regulation is mediated in part by RORA, being as a potential target for anti-fibrogenic therapy. We believe that understanding the metabolic adaptations of the dHSCs trajectory could be a novel strategy to provide new and potentially targeted interventions.

The current study highlights the significance of examining cell trajectories during differentiation and disease. It also shows that the mechanisms controlling human development and fibrosis can be recapitulated using iPSC differentiation models, which enables the identification of crucial transcription factors for cell commitment and disease.

## Acknowledgments

P.S-B. is supported *by Instituto de Salud Carlos III (ISCIII) “*FIS PI20/00765”, “DTS22/00032”, AGAUR “2019 PROD 00055”, “2021 PROD 00035” and co-founded by the European Union “HORIZON-HLTH-2022-STAYHLTH-02-01”. R.A M-G is funded by *Instituto de Salud Carlos III* “FI18/00215”, S.A. received a grant from the *Ministerio de Educación, Cultura y Deporte* “FPU17/04992”, C.M is funded by *Centro de Investigación en Red Enfermedades Hepáticas y Digestivas (CIBERehd)*. S.A. is funded by la Caixa Foundation “100010434” and from the European Union’s Horizon 2020 under the Marie Skłodowska-Curie “847648” and PID2021-124694OA-I00 from MCIN/AEI/10.13039/501100011033 and *FEDER Una manera de hacer Europa*. A.G. is supported by Competitiveness (MINECO PID2020-15591RB-100), La Marató de TV3 (202001-32) and *CERCA Programme/ Generalitat de Catalunya* for institutional support. J.P. is supported by *La Marató de TV3* (202001-32). D.R-M. is supported by *Deutsche José Carreras Leukämie-Stiftung* (DJCLS 13R/2022).

## Authors contribution

R.A.M.G.T. conceived the study, designed and performed the experiments, interpreted the results and wrote the manuscript. J.V., Q.X., S.A.M., B.A-B., P.R.B., M.F-F., A.N-G., A.B-R., P.S.F, J.P.G, D.R-M., P.A.G., C.M.S., L.Z., L.S., P.C.V., M.V., A.W., A.M., performed experiments. M.A. and F.E. performed the proteomic analysis. J.J.L. performed computational analysis. B.A., M.M-C., A.G., A.P., A.Z., I.G., M.C., S.A., provided insights. P.S-B., conceived the study, interpreted the results, supervised the experiments, and wrote the manuscript.

## STAR METHODS

### EXPERIMENTAL MODELS AND SUBJECT DETAILS

#### iPSCs cell line

For the proteome characterization and all the subsequent *in vitro* assays, we have used the human KULi003-A iPSC line (BioSamples Cat#SAMEA110177409).

#### HepG2 cell line

The HepG2 cell line was purchased from Sigma (Cat#8511430).

#### Staggerer mice

All animal experiments were approved by the Ethics Committee of Animal Experimentation of the University of Barcelona and were conducted in accordance with the National Institute of Health Guide for the Care and Use of Laboratory Animals. For the fibrogenic experiments in staggerer mice (sg/sg), mice homozygous for the staggerer spontaneous mutation, both genders of between 8-10 weeks of age were used.

#### WT mice

All animal experiments were approved by the Ethics Committee of Animal Experimentation of the University of Barcelona and were conducted in accordance with the National Institute of Health Guide for the Care and Use of Laboratory Animals. For the fibrogenic experiments, C57BL/6J female and male mice of 8-10 weeks of age were used.

## METHOD DETAILS

### iPSCs differentiation towards dHSCs

iPSCs KULi003-A, clone 5, clone 8 and clone 13J of RORA^+/-^ iPSCs were differentiated following the protocol described previously^34^. Briefly, dHSCs differentiation started when iPSCs achieved 70% confluence. From day 0 to day 4, cells were cultured in liver differentiation media (LDM) supplemented with BM4 (20ng/ml, R&D, Cat#314-BP-010). From day 4 to day 6, cells were treated with BMP4 (20ng/ml), FGF1(20ng/ml, R&D, Cat#232-FA-025) and FGF3 (20ng/ml, R&D, Cat#1206-F3-025). From day 6 to day 8, cells were stimulated with FGF1 (20ng/ml), FGF3 (20ng/ml), retinol (5μM, Sigma Aldrich, Cat#R7632-100MG) and palmitic acid (100μM, Sigma Aldrich, Cat#P0500). Finally, from day 8 to day 12, cells were cultured in LDM supplemented with retinol and palmitic acid.

### Generation of heterozygote KO iPSC for RORA gene

Heterozygous RORA-knockout (KO) iPSCs (RORA^+/-^) were obtained as described elsewhere^35^. The sgRNA “5’-ATCTGTGGAGACAAATCATCAGG-3’” (IDT) was designed by the CRISPR tool (https://bioinfogp.cnb.csic.es/tools/breakingcas). Rock inhibitor at 10μM (Y-27632, BD Bioscience, Cat# 562822) was added to the iPSCs 3 hours (h) before nucleofection. 100 pmol Alt-R™ S.p. HiFi Cas9 Nuclease V3 (IDT) was incubated with 120 pmol Alt-R® CRISPR-Cas9 sgRNA (IDT) at 25°C for 10 minutes (min). 200.000 cells were dissociated with Accutase (Gibco, Cat#A1110501), washed twice with PBS without Ca and Mg and resuspended with 20μl of P3/S1 Buffer. RNP complex was added to the cells prior to nucleofection in a 20ul cuvette. Cells were nucleofected with the 4-D Nucleofector System (Lonza, Cat#AAF-1003X) using the CA-137 program. Nucleofected cells were cultured in a 12 well plate, with mTSR1 and 10uM of Y-27632. After 72 h of recovery, 1000 cells were seeded at a single cell level in a 100 mm plate to generate single-cell colony. Genotyping was performed by PCR and Sanger sequencing from single cell colonies to analyze the genotype. Two rounds of single cell cloning were performed to ensure a single cell clone.

### Chemical modulation of RORA signalling

On day 2 of the differentiation protocol, the synthetic agonist (SR1078, Merck, Cat#557352) or antagonist (SR1001, MCE, Cat#HY-13421) of RORA and RORG was added to the cells at 0.1mM or 85nM, respectively. Treatment refresh was performed every two days. Samples were collected at day 12 for further analysis. For the modulation of RORA after passage, differentiated cells at day 12 were passaged as explained previously ^14^. Once cells achieved 70% confluence, SR1001 (85nM) and SR1078 (3mM) were added for 24 hours. Samples were collected for further analysis.

### Fibrogenic *in vitro* assay

dHSCs were passaged and treated with TGFβ (Prepotech, Cat#AF-100-21C) at 5ng/mL for 24h or 7 days as explained elsewhere^17^ . During the last 24h, SR1078 (3mM) was added to the cell culture and samples were collected fur further analysis.

### Co-culture 3D assay

Liver spheroids were generated using dHSCs and HepG2 in a 1:2 ratio. 3000 cells were seeded in a non-adherent U-Shape bottom 96-well plate in DMEM Glutamax, 10% FBS and 1% P/S. Plates were centrifuge at 200g for 2 min. and placed on the incubator. After 3 days, a half medium change was performed, and experiments started after 5 days of culture. Then, liver spheroids were treated with SR1078 at 3mM for 24h; 10ng/mL for TGFβ during 48h. During the last 24 hours, liver spheroids were also treated with SR1078 3mM. Samples were collected for further analysis.

### In vitro studies with rat neonatal cardiomyocytes (NCMs) and fibroblasts (NCFs)

Rat NCMs were obtained from the ventricles of 1-to 3-day-old Sprague-Dawley rats. Briefly, hearts were digested with a collagenase solution (Collagenase Type I; Life Technologies, Cat# 17100017) and differential plating was performed. NCMs were plated at a density of 0.5 × 106 cells/well on 6-well culture plates and grown for 24 h. NCMs were treated with phenylephrine PE (10 μmol/L; Sigma, Cat#P1240000), as a pathological hypertrophic growth factor for 24 h. Rat neonatal cardiac fibroblasts (NCFs) were isolated from the ventricles of 1-to 2-day-old Sprague-Dawley rats. Passage 1 and 3 cells were plated at a density of 0.3 x 105 cells/cm2 on culture dishes and grown for 24 h.

### Sample preparation for proteome analysis

Day 0, day 2, day 4, day 6, day 8, day 10, day 12 and pHSCs were washed once with DPBS -/- and 500mL of buffer lysis was added to the culture (7M urea, 2 M thiourea and CHAPS 4%) following the filter aided FASP protocol described elsewhere _22_ with minor modifications. Trypsin was added at a trypsin:protein ratio of 1:50, and the mixture was incubated overnight at 37°C, dried out in a RVC2 25 speedvac concentrator (Christ), and resuspended in 0.1% formic acid. Peptides were desalted and resuspended in 0.1% FA using C18 stage tips (Millipore).

### Mass spectrometry analysis

Samples were analyzed in a hybrid trapped ion mobility spectrometry – quadrupole time of flight mass spectrometer (timsTOF Pro with PASEF, Bruker Daltonics) coupled online to a nanoElute liquid chromatograph (Bruker). The sample (200ng) was directly loaded in a 15 cm Bruker nanoelute FIFTEEN C18 analytical column (Bruker) and resolved at 400 nl/min with a 30 min gradient. The column was heated to 50°C using an oven. Protein identification and quantification was carried out with MaxQuant software using default settings^24^. Searches were performed against a database consisting of human protein entries (Uniprot/Swissprot), with precursor and fragment tolerances of 20ppm and 0.05 Da. Only proteins identified with at least two peptides at FDR<1% were considered for further analysis. Data (LFQ intensities) was loaded onto a Perseus platform 25 and further processed (log2 transformation, imputation). Protein abundances were normalized against the day 0 for each individual.

### RNA extraction, cDNA synthesis, qRT-PCR analysis and Bulk RNA sequencing

RNA from cells was obtained using the Quick RNA-Microprep Kit (Zymo, Cat#R1050), following the manufacturer’s instructions. For 3D cultures, total RNA from 6 pooled spheroids were extracted using the same kit. Total RNA was extracted from mouse whole liver tissue using Trizol (Life Technologies, Carlsbad, CA, Cat#15596026). The total RNA extracted underwent quality control by Bioanalyzer 2100 (Agilent Technologies, Santa Clara, CA). RNA concentrations were measured using the Nanodrop 1000 (ThermoFisher). cDNA synthesis was performed with the MultiScrib (Applied biosystems, Cat#4368813) followed by a quantitative reverse transcription polymerase chain reaction (qRT-PCR) on an ABI 7900HT cycler (Applied Biosystems) and SYBR green master mix (Life Technologies, Cat#11733046). Gene expression was normalized to *GAPDH* for *in vitro* systems and *Actb* for in vivo experiments. Expression values were calculated based on the ΔΔCt method. For bulk-RNA sequencing, the quality of RNA was evaluated using Agilent RNA 6000 Nano Chips (Agilent Technologies, Santa Clara, CA, USA). Sequencing libraries were prepared following the TruSeq Stranded mRNA Sample Preparation with the corresponding kit starting from 125 ng of total RNA per sample. Sequencing of the mRNA libraries was performed in a HiSeq2500 (Illumina Inc, San Diego, CA, USA) to obtain at least 30 million 51nt-single-end reads per sample. GO biological processes term enrichemnet, KEGG pathway, Reactome and gene set enrichment analysis were performed using DAVID ^36^, and GSEA database^37^.

### Intracellular glucose level testing

Intracellular glucose levels from day 4 differentiated cells and dHSCs passaged cells were assessed using the Glucose-Glo™ Assay, following manufacturer instructions (Promega, Cat#TM494). Briefly, the assay was applied to cells in 96-well plate. Before beginning the assay, the medium was removed, and the cells were washed with 100 μl of DPBS. To initiate glucose uptake, 50 μl of 2DG in DPBS were used. The uptake reaction was stopped, and the samples were processed following the manufacturer’s instructions.

### XF24 extracellular flux

The oxygen consumption ratio (OCR) was determined using Seahorse XF Cell Mito Stress Test (Agilent, Santa Clara, CA, US). Briefly, iPSC-RORA +/- and their WT counterparts, treated and untreated with the RORA agonist SR1078 at 0.1mM at day 4 of the differentiation, were sequentially treated with 1 μM oligomycin; 2 μM carbonyl cyanide m-chlorophenyl hydrazone (CCCP); and a 1 μM mixture including rotenone and antimycin A according to manufacturer’s instructions. Seahorse XFe Wave Software (Agilent) was applied to analyze the data. Similarly, cells after passage were treated with RORA agonist and/or TGFb for 24h. Finally, the cell mass was normalized quantifieng total protein using the BCA protein assay kit (ThermoFisher, Cat# 23225).

### Immunofluorescence and immunohistochemistry

Cells were fixed using 4% Neutral Buffered Formalin for 15 min and washed with PBS for three times before permeabilization at room temperature during 30 min. with 0.2% Triton X-100 in PBS. Afterwards, the cells were blocked with 3% of donkey serum at room temperature. Cells were incubated overnight at 4°C with primary antibodies in DAKO REALTM Anti-body diluent (Agilent, Cat#S2022). Secondary antibodies (1:500 dilution) were incubated for 30 min. To ensure nuclear staining, samples were mounted with Mounting Medium with 4’,6-diamidino-2-phenylindole (DAPI) (Vector Laboratories, Cat# H-1200-10). Paraffin embedded liver sections (3 μm) from human and mouse liver samples were stained for aSMA (1:100, Abcam, Cat#ab5694), KI67 (1:50, Abcam, Cat#ab16667), MPO (1:50, Abcam, Cat#ab9535), F4/80 (1:200, BioRad, Cat#MCA497) and CD3 (1:750, BioRad, Cat#MCA1477). Sections were deparaffinized and incubated in Target Retrieval Solution, citrate pH6 (Dako, Cat#2369), heated in a pressure cooker for 20 min. or rehydrated and antigen retrieved with EnVision Flex Target Retrieval Solution Low or High pH (Agilent, Cat#K8004 for the high and Cat#K8005 for the low pH) in a Dako PT Link. Samples were incubated with primary antibody overnight at 4°C. After PBS washes, sections were incubated with secondary antibody. Diaminobenzidine (Agilent, Cat#K346811-2) was used as a chromogen and finally, sections were counterstained with hematoxylin. For immunofluorescence staining, after incubation with secondary antibody, sections were mounted with Mounting Medium for Fluorescence with 4’,6-diamidino-2-phenylindole for nuclear staining. The antibodies used are shown in Resource table.

### RORA immunofluorescence in human tissue

Human liver sections were included in OCT and maintained at -80°C. Slides were fixed at 4% PFA for 2 min. They were then washed in PBS for a few seconds and blocked in 100µL 5% BSA in PBS + 0,3% TritonX100 per tissue section during 1h. Tissue sections were incubated over night with primary antibodies (RORA 1:50, Abcam, Cat#ab70061) in 5%BSA/PBS/Triton at 4°C in dark moist chamber. Then, the sections were incubated 1h with secondary antibodies in 5%BSA/PBS/Triton at room temperature (RT) in a dark moist chamber. Tissue sections were mounted using the Vectashield Mounting medium for fluorescence with DAPI.

### In vivo test in carbon tetrachloride induced liver fibrosis model in homozygous staggerer mice

To test the physiological role of RORA upon liver fibrosis in vivo, we established a carbon tetrachloride (CCl4) model based on intraperitoneal (I.P.) injection of 0,5ml/kg of CCl4 diluted in corn oil. Mice were injected two days a week during four 4 weeks. Blood and liver tissues were collected for further analysis.

### *In vivo* test of RORA as an antifibrogenic target in fibrogenic models

For the liver fibrosis model, 14 C57BL/6J mice were injected twice a week with CCl4 0,5ml/kg (25% corn oil) for 4 weeks. During the last 2 weeks, mice were divided into two groups: 1) treated with SR1078 (10mg/kg) by I.P. injections twice a week; 2) vehicle treated mice received I.P. injections twice a week. SR1078 was diluted in 5% DMSO to a final concentration of 2mg/ml. For the heart fibrotic model, 10 C57BL/6J mice received isoproterenol (Sigma Aldrich, Cat#SLC62971) administered by continuous infusion of 60mg/kg/day using minipumps (Model 1007D, Alzet). Animals were divided into two groups: 1) treated with SR1078 (10mg/kg) every 48h and 2) vehicle treated mice for 1 week. Animals were sacrificed after one week. For the kidney fibrotic model, 19 C57BL/6J mice underwent ligation of the left ureter and the mice were divided into two groups: 1) treated with SR1078 (10mg/kg) from day 3 after surgery until the end; 2) vehicle treated mice from day 3 after surgery for 15 days. Blood and liver, heart and kidney tissues were collected for further analysis.

### Collagen tissue determination

The amount of collagen in liver tissues was determined by measuring the content of hydroxyproline using a Hydroxyproline assay kit (Abcam, Cat#MAK008-1KT) as described by the manufacturer. Absorbance was measured at 540 nm using a FLUOstar OPTIMA FL reader (BMG LABTECH).

### Biochemical determination by Echevarne laboratories (Barcelona, Spain)

Blood from mice included in the experimental studies was collected and analyzed for alanine transaminase (ALT), aspartate transaminase (AST), phosphatase alkaline (AP) and lactate dehydrogenase (LDH) values.

### Statistical analysis

GraphPad Prism v8.0 was used for the statistical analyses. The D’Agostino-Pearson omnibus normality test, Anderson-Darling test and Shapiro-Wilk normality test were performed to assess data distribution. For statistical analysis of parametric data, the two-tailed unpaired Student’s t test was used for groups of two; one-way ANOVA followed by Sidak multiple comparison posthoc tests were used for comparison of more than two groups. For non-parametric data, the Mann-Whitney U test was used for groups of two while the Kruskal-Wallis test followed by Dunn multiple comparison posthoc test was used for comparison of more than two groups.

---

**Table S1** is related to figure 1 and comprehends proteins included in each cluster, according to their changes along differentiation and the gene ontologies (GO) enriched in each cluster of proteins (table S1B – GO enriched in matrisome cluster; table S1C – GO enriched in metab cluster; table S1D – GO enriched in pluri cluster; table S1E – GO enriched in proli cluster; table S1F – GO enriched in basal cluster).

**Table S2** is related to figure 2 and contains the TFs predicted to regulate the transition between phases. Table S2A are the TFs predicted to regulate the transition between phase 1 and phase 2. Table S2B are the TFs predicted to regulate the transition between phase 2 and phase 3. Table S2C are the TFs predicted to regulate the transition between phase 3 and pHSCs.

**Table S3** is related to figure 3 and contains the transcriptional analysis of the cells treated with SR1078 during differentiation and after passage. Table S3A contains the gene ontologies of cells treated with SR1078 during differentiation. Table S3B contains the reactome pathways enriched with SR1078 during differentiation. Table S3C contains the gene onotologies of cells treated with SR1078 after passage. Table S3D contains the reactome pathways enriched with SR1078 after passage.

## Supplementary Figure Lengends

**Figure S1: Protein clustering of the proteome profile of dHSCs differentiation and comparison with primary HSCs.** (A) Heatmap showing protein clustering according to protein dynamics across differentiation. (B) TOP 5 gene ontologies (GOs) enriched in each cluster identified across differentiation.

**Figure S2: RAR-related Orphan Receptor A is a potential driver of the dHSCs phenotype by improving mesoderm commitment and modulating the dHSCs activation state.** (A) Representative microscopy images of dHSCs treated along differentiation with the RORA agonist at 0,1mM. (B) Gene expression of treated cells across differentiation with SR1078 of activation markers (*ACTA2*, *COL1A1* and *LOX*) upon TGFb insult (10ng/ul) during 24h. (C) Sanger sequencing of the iPSC-RORA (+/-). (D) Western Blot for RORA of BJ1 iPSCs wild type (WT) cell line and three different clones. (E) immunofluorescence for pluripotency marker (OCT4) and RORA in three iPSC-RORA (+/-) clones and wildtype (WT). (F) Gene expression of pluripotency markers in three three iPSC-RORA (+/-) clones and wildtype (WT). (G) Karyotype of the three iPSC-RORA (+/-) clones. (H) Schematic representation of the experimental design for the treatment during the differentiation of iPSCs towards HSCs with RORA invers agonist (SR1001). (I) Representative images at day 12 of Sirius Red staining in the cell culture of untreated (Vehicle) and treated (SR1001) cells across differentiation. (J) Gene expression of quiescent (*LHX2*, *RELN*) and activation (*ACTA2*, *COL1A1*) markers after SR1001 treatment. (K) Pathway enrichment analysis show that passage dHSCs are activated cells. (L) Passaged dHSCs treated with SR1001 during 24h. (M) Gene expression of activation markers (*COL1A1, ACTA2, LOX*) treated with the RORA antagonist SR1001 during 24h at passage. Significant differences are indicated as *p<0.05.

**Figure S3: RORA regulates the activation of HSCs, leading to the progression of liver fibrosis.** (A) Hydroxyproline assessment in basal livers from staggerer mice (sg/sg) and their wildtype (wt) counterparts. (B) Gene expression of activated HSCs markers (*Acta2*, *Col1a1*, *Timp1* and *Mmp2*) in basal livers from staggerer mice (sg/sg) and their wildtype (wt) counterparts. (C) Alanine aminotransferase (ALT), aspartate aminotransferase (AST), lactate dehydrogenase (LDH) and alkaline phosphatase (AP) determination in staggerer mice (sg/sg) and their wildtype (wt) counterparts after 4-week treatment with CCl_4_. (D) Immunofluorescence for macrophage (F4/80), neutrophils (MPO), lymphocytes (CD3), and proliferation (KI67) determination after CCl4 treatment in sg/sg and wt mice.

**Figure S4: RORA agonist reduces fibrosis**. (A) Gene expression of *Mmp2*, *Mmp9*, *Mmp12*, *Il1b*, *Il6* and *Tnfa* in CCl_4_ fibrosis model treated with the RORA agonist (SR1078) for 2 weeks. (B) Alanine aminotransferase (ALT), aspartate aminotransferase (AST), lactate dehydrogenase (LDH) and alkaline phosphatase (PA) determination in CCl_4_ fibrosis model treated with the RORA agonist (SR1078) for 2 weeks. (C) Immunofluorescence for macrophage (F4/80), neutrophils (MPO), lymphocytes (CD3), and proliferation (KI67) determination in CCl_4_ fibrosis model treated with RORA agonist (SR1078) for 2 weeks. (D) Illustration of the experimental design of unilateral uteral obstruction (UUO) intervention. (E) Gene expression of RORA in the damaged kidney (left kidney - LK) and the control kidney (right kidney - RK). (F) Representative image of the Sirius red and αSMA staining of the left kidneys from vehicle and SR1078 treated groups. (G) Gene expression of *Rora*, *Acta2* and *Col1a1* of left kidneys from vehicle and SR1078 treated groups.

