## Supplementary figures and images for "Trajectory Analysis of Hepatic Stellate Cell Differentiation Reveals Metabolic Regulation of Cell Commitment and Fibrosis"

### Supporting information and supplementary figures

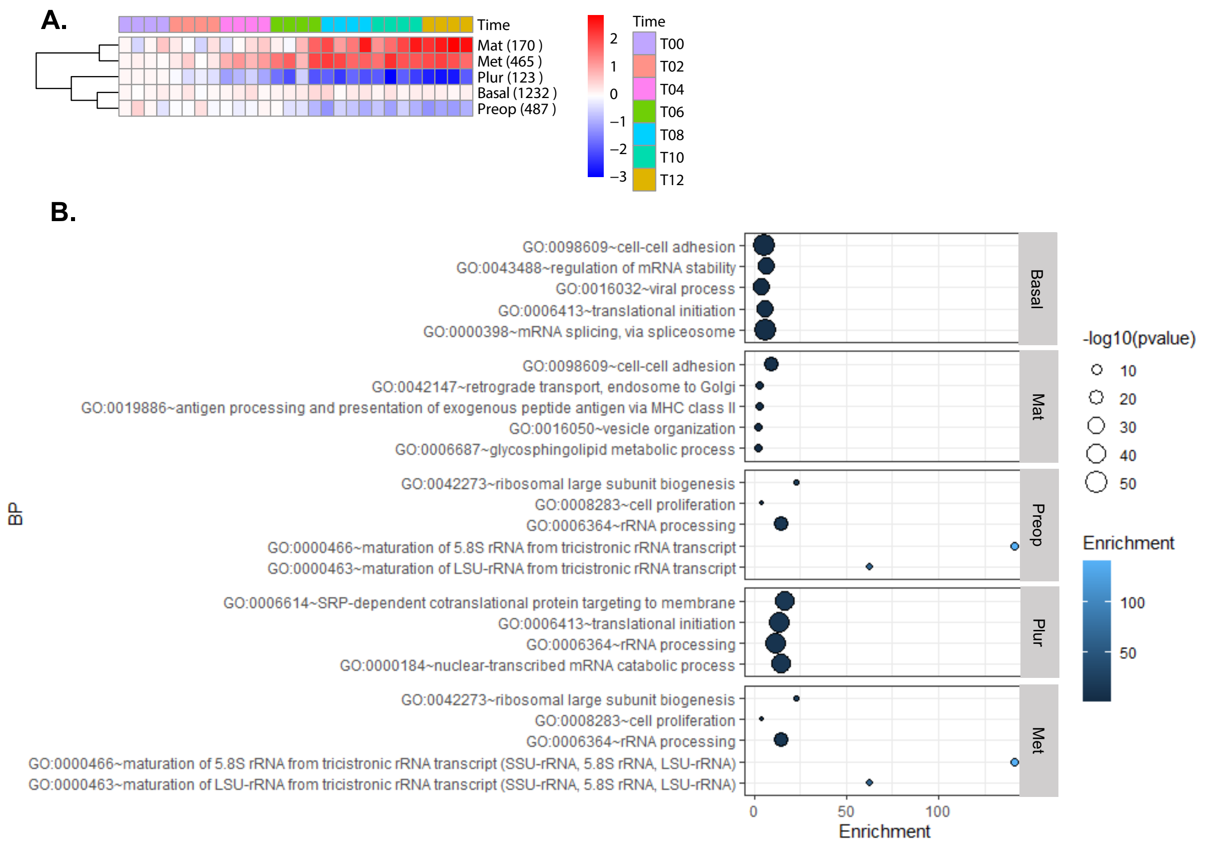

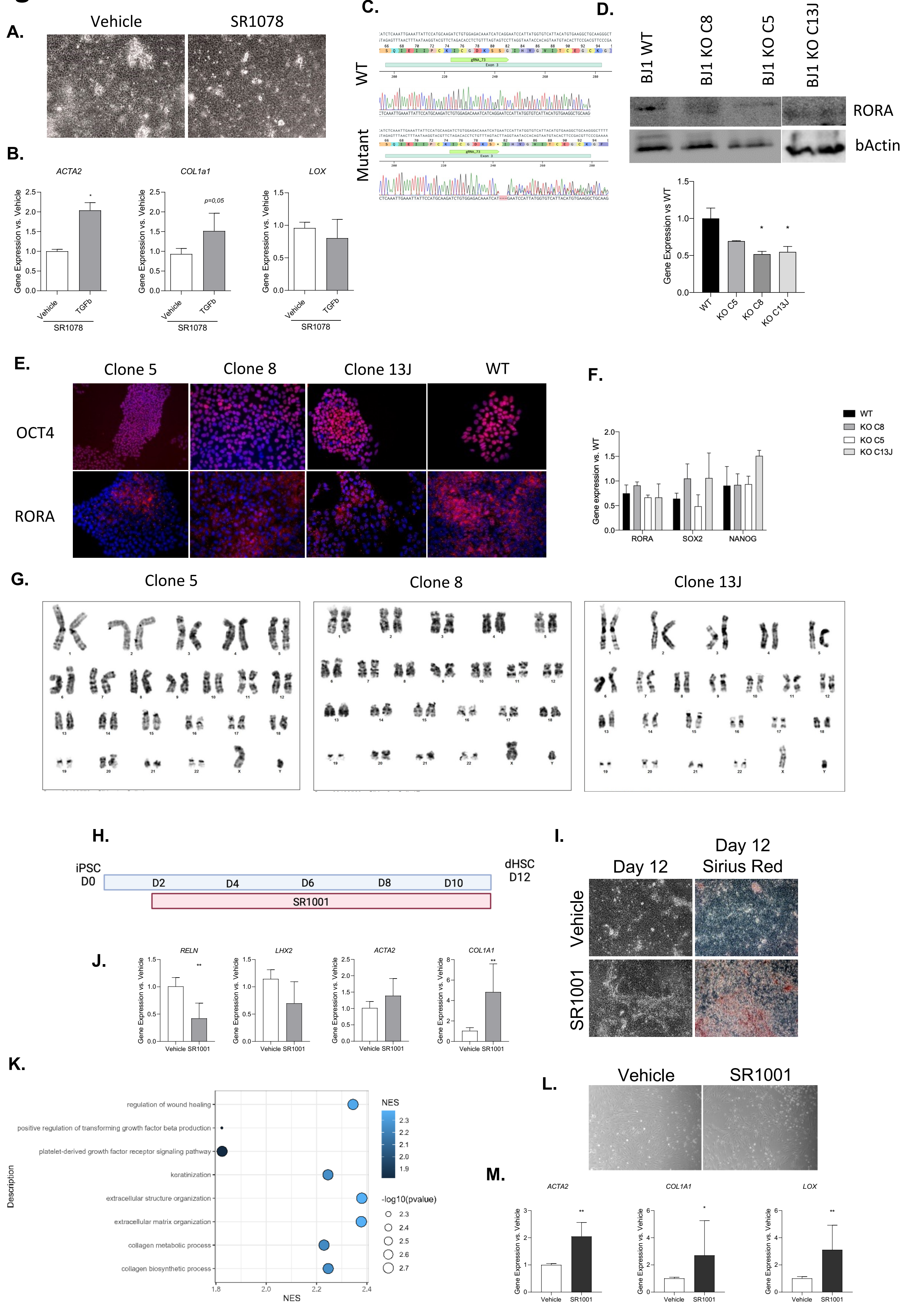

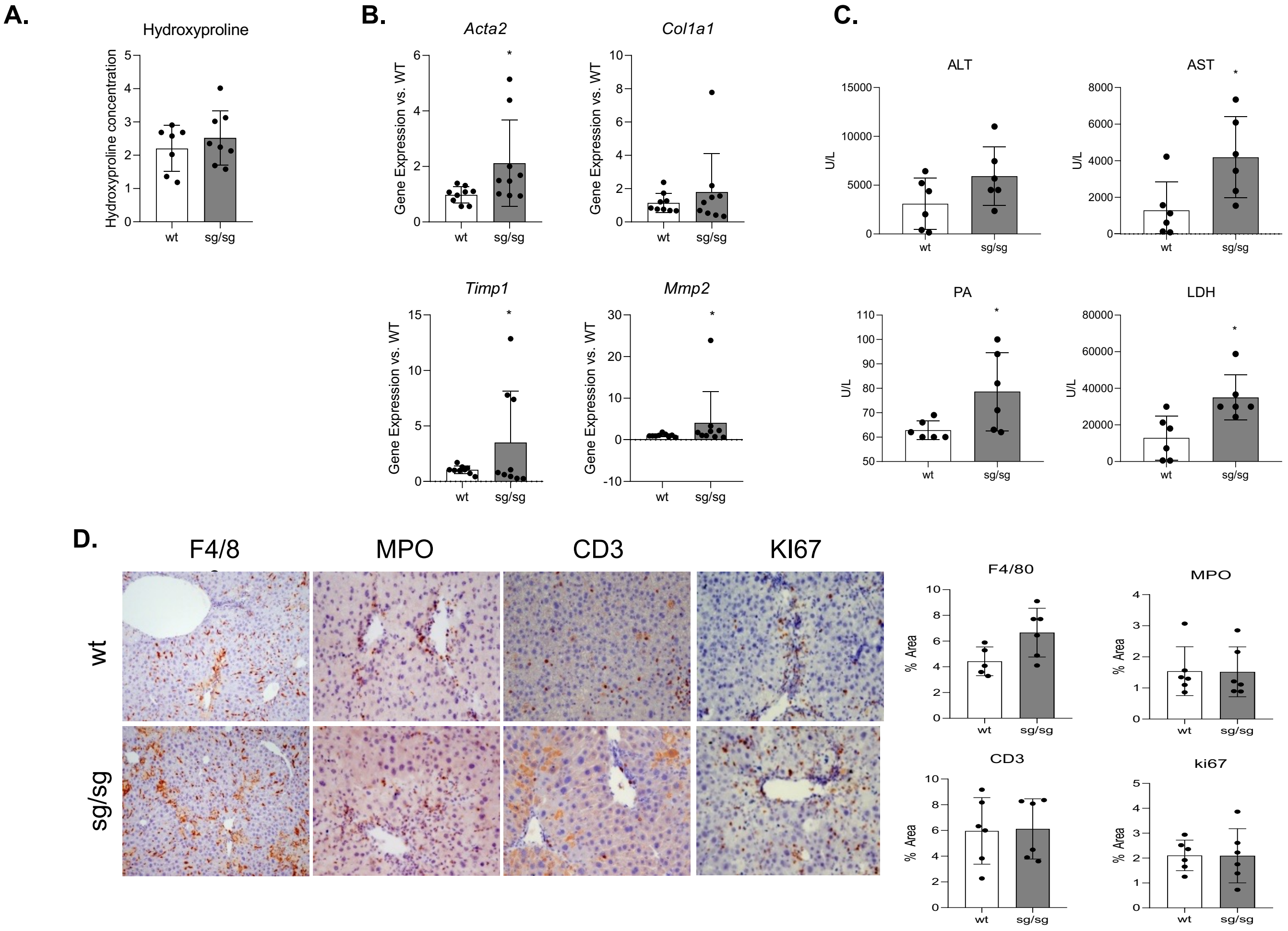

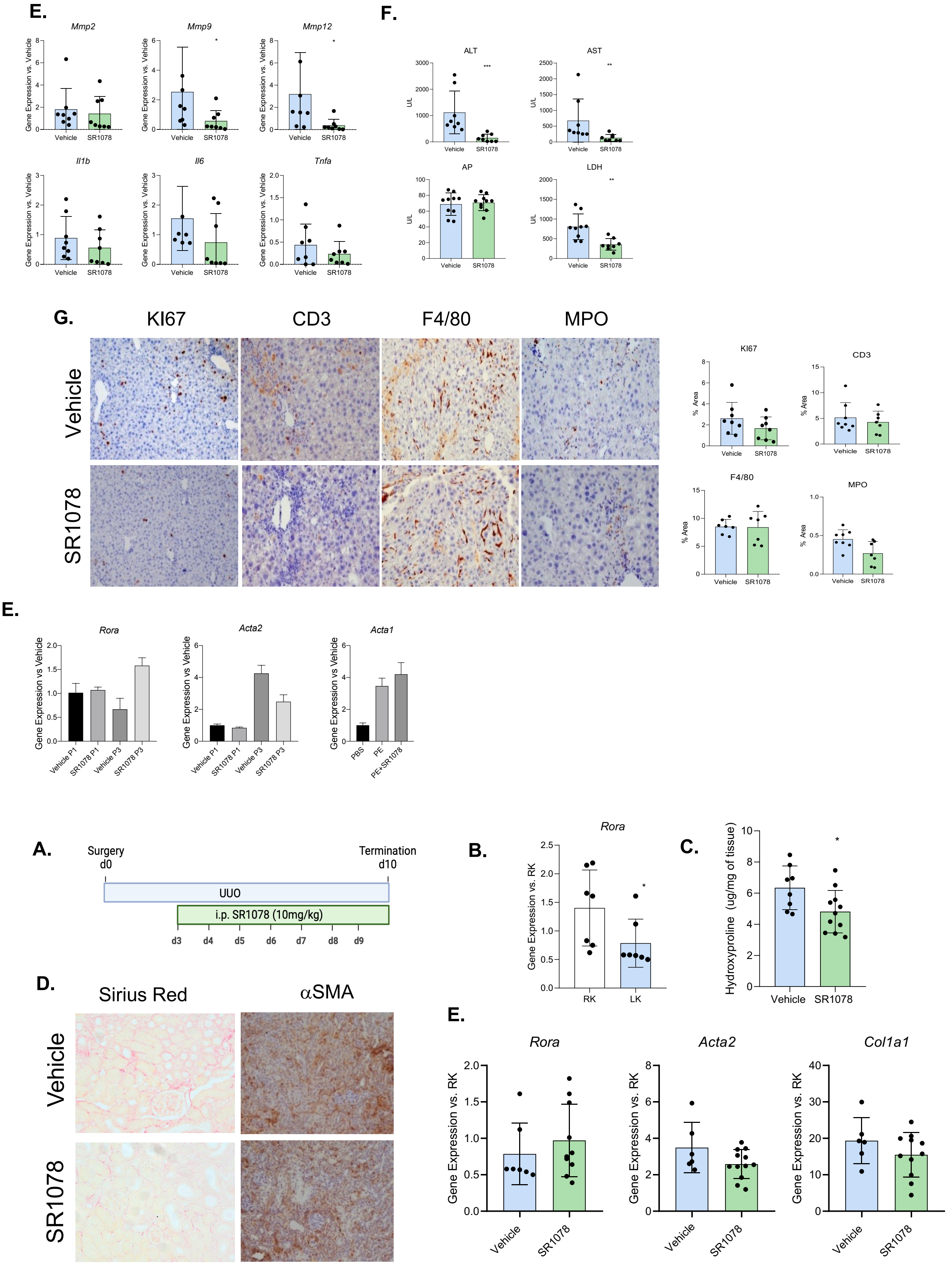
